# Tuning gene expression variability and multi-gene regulation by dynamic transcription factor control

**DOI:** 10.1101/287565

**Authors:** Dirk Benzinger, Mustafa Khammash

**Affiliations:** Department of Biosystems Science and Engineering (D-BSSE), ETH–Zürich, Mattenstrasse 26, 4058 Basel, Switzerland

## Abstract

Many natural transcription factors are regulated in a pulsatile fashion, but it remains unknown whether synthetic gene expression systems can benefit from such dynamic regulation. Using a fast-acting, light-responsive transcription factor in *Saccharomyces cerevisiae*, we show that dynamic pulsatile signals reduce cell-to-cell variability in gene expression. We then show that by encoding such signals into a single input, expression mean and variability can be precisely and independently tuned. Further, we construct a light-responsive promoter library and demonstrate how pulsatile signaling also enables graded multi-gene regulation at fixed expression ratios, despite differences in promoter dose-response characteristics. Pulsatile regulation can thus lead to highly beneficial functional behaviors in synthetic biological systems, which previously required laborious optimization of genetic parts or complex construction of synthetic gene networks.

## INTRODUCTION

The relationship between gene expression and cellular phenotype lies at the center of many questions in different branches of biological research. While strong perturbations of gene expression like knock-outs and overexpression led to a tremendous increase in our understanding of protein function, graded gene expression regulation allows us to obtain a quantitative understanding of the expression-phenotype mapping. Furthermore, conditional and titratable gene expression is of major importance in biotechnology and synthetic biology. Thus, a variety of tools for regulating cellular protein levels, such as gene expression systems based on hormone or light-inducible transcription factors, were developed ^1^. With a few exceptions ^2–4^, expression levels are regulated by adjusting the strength of an input, leading to a graded and constant activation of a transcriptional regulator (**Fig. 1a**, from here on referred to as amplitude modulation (AM)). In contrast, recent studies have shown that many natural regulatory proteins, including transcription factors (TFs), exhibit pulsatile activation patterns leading to a variety of phenotypic consequences ^5^.

**Figure 1.**
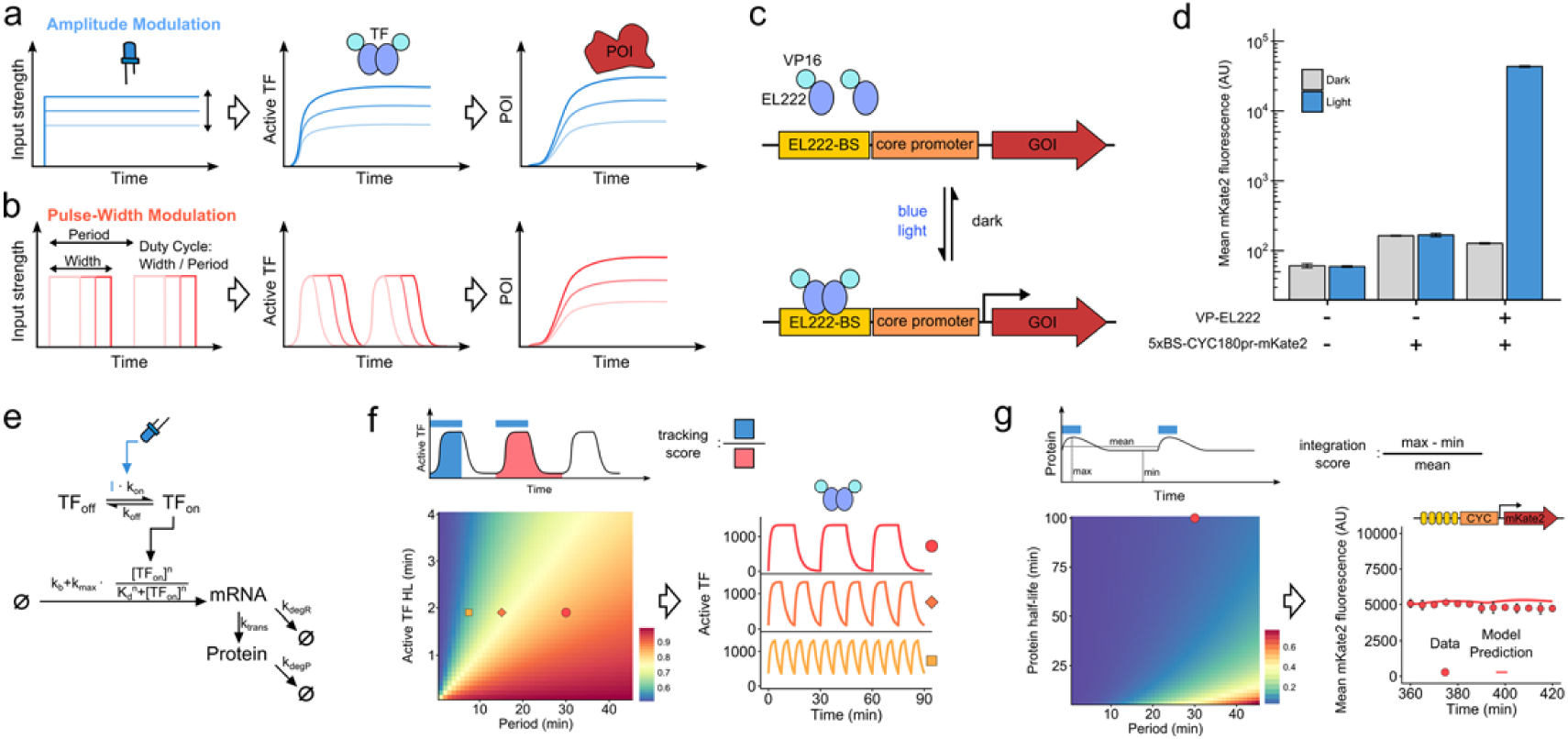
Characterization of an EL222 based optogenetic gene expression system in *S. cerevisiae*. **(a)** and **(b)** Schematic of gene expression regulation by AM (a) and PWM (b). Input signals (left) lead to TF activation (middle) and finally expression of a protein of interest (POI, right). **(c)** Illustration of the optogenetic gene expression system. Blue light triggers structural changes in VP-EL222 leading to dimerization, binding to its cognate binding site in a synthetic promoter region, and finally transcription of a gene of interest (GOI). **(d)** Effect of blue light illumination on VP-EL222 mediated gene expression. Strains, with or without the VP-EL222 and a reporter construct (5xBS-CYC180pr-mKate2), were grown either in the dark or under illumination (460 nm LED source, 350 µW/cm^2^) for 6 h. The average cellular mKate2 fluorescence was measured using flow cytometry. Data represents the mean and s.d. of three independent experiments. (**e**) Graphical representation of the model describing VP-EL222 mediated gene expression. The model consists of three ordinary differential equations describing VP-EL222 / TF activity, TF-mediated mRNA production with a transcription rate modeled by Hill kinetics, and protein expression. The light input is denoted by **I**. Arrows depict reactions (inferred parameter values are shown in Supplementary Table 1). See Methods and Supplementary Note 1 for ordinary differential equations and further details on the modeling. (**f**) Model-based analysis of pulsatile TF behavior upon PWM. To quantify the temporal TF response, we use a tracking score defined by the ratio between the integrated TF activity during the light pulse and the whole period (top). The heatmap depicts the tracking score for a 50% duty cycle and a simulated light intensity of 420 µW/cm^2^ as a function of the PWM period and the half-life (HL) of the active VP-EL222 state. Three PWM induction regimes that are used throughout this study are marked on the heatmap. On the right, predicted temporal TF activities are shown for these conditions. (**g**) Model-based analysis of PWM-mediated protein expression. Ideally, PWM should not result in significant temporal variations of protein levels. To quantify this behavior, we use a score defined by the ratio of the maximal expression difference during the period divided by the mean expression level (top). The heatmap depicts the score for a 10% duty cycle as a function of the PWM period and protein half-life. On the right, the predicted and measured time-course of FP expression in response to two successive light pulses with a 30 min period are shown after 360 min of induction at 10% duty cycle. Data represents the mean and s.d. of two independent experiments.

Motivated by the occurrence of pulsatile transcription factor regulation in natural systems, we hypothesized that synthetic gene expression systems can benefit from such dynamic regulation. To test this hypothesis, we constructed a fast-acting, and genomically integrated, optogenetic gene expression system based on the bacterial light-oxygen-voltage protein EL222 in *Saccharomyces cerevisiae* ^4^. Fast kinetics of the optogenetic TF together with the ability to control light intensity with high temporal precision allowed us to tune gene expression using pulsatile TF inputs. In particular, we performed pulse-width modulation (PWM) ^3^, meaning that the duration of input pulses is varied to achieve different gene expression levels, while keeping the period of the pulses constant (**Fig. 1b)**. The ratio of pulse duration to the period is referred to as duty cycle. PWM can be performed at different input amplitudes and periods, providing further options for dynamic signal encoding to regulate gene expression levels. We used a mathematical model to identify suitable PWM periods and then showed experimentally that these can be exploited to tune gene expression properties. By comparing this PWM approach to AM, we establish that dynamic encoding of pulsatile signals can drastically increase the functionality of gene expression systems.

## RESULTS

### Characterization and modeling of an EL222-based expression system in *S. cerevisiae*

In order to regulate gene expression using PWM, we implemented an optogenetic gene expression system based on a previously described TF consisting of a nuclear localization signal, the VP16 activation domain ^6^, and the light-oxygen-voltage domain protein EL222 of *Erythrobacter litoralis* (VP-EL222) ^4^. Blue light illumination triggers structural changes in EL222 leading to homodimerization and binding to its cognate binding site (**Fig. 1c**). An EL222-responsive promoter was constructed by inserting five binding sites for EL222 ^4^ upstream of a truncated CYC1 promoter (5xBS-CYC180pr) and was used to drive the expression of the fluorescent protein (FP) mKate2 ^7^. For initial characterization, we measured the expression levels of mKate2 in the dark and after 6h of blue light illumination via flow cytometry (**Fig. 1d**). Illumination led to a VP-EL222 dependent increase in cellular fluorescence of more than 250-fold. In the dark, the presence of VP-EL222 did not affect gene expression. Neither the expression of VP-EL222 nor light-induction affected cell growth or constitutive gene expression (**Supplementary Fig. 1**).

In order to achieve a quantitative understanding of the system and investigate potential PWM regimes, we derived a simple mathematical model of VP-EL222 mediated gene expression (**Fig. 1E**, for details see **Supplementary Note 1**). The model was fitted to the data of three characterization experiments, namely a gene expression time-course as well as dose response curves to AM and PWM with a period of 7.5 min (**Supplementary Fig. 2**). Analyzing the model showed the importance of fast VP-EL222 deactivation kinetics for successful PWM (**Supplementary Note 1.3**). For a fixed pulse width, slow deactivation rates require long PWM periods to achieve purely pulsatile TF regulation (**Fig. 1f**). However, such periods may result in significant temporal variation of downstream gene expression / input tracking (**Fig. 1g**). Here, the half-life of the active VP-EL222 state was inferred to be lower than 2 minutes (**Fig. 1f**). Measurements of transcription upon a blue light pulse using smFISH lead to results consistent with the fast VP-EL222 kinetics (**Supplementary Fig. 3, Supplementary Note 1.4**). For the inferred deactivation rate, the model predicts strongly pulsatile TF activity for a 30 min PWM period and a 50% duty-cycle, whereas for the same duty-cycle TF activity does not return to baseline when a 7.5 min PWM period is used (**Fig. 1f**). Importantly, even for a 30 min period, temporal changes in protein expression at steady state are expected to be minor for a wide range of protein half-lives (**Fig. 1g, Supplementary Note 1.3**). We confirmed experimentally that there is no measurable input tracking for a stable fluorescent protein (**Fig. 1g**). Thus, the fast kinetics of the VP-EL222 based system together with its tight regulation, and apparent lack of toxicity, makes it an ideal gene expression tool for a variety of applications and enables the regulation of protein levels by PWM.

### Pulsatile input signals achieve coordinated multi-gene expression

Given that most cellular phenotypes are a result of the coordinated regulation of many genes whose protein expression ratios can be of high importance for achieving these phenotypes ^8^, we explored the use of AM and PWM for achieving graded expression of multiple proteins, each at a different level, with a single gene expression system. Eukaryotic genes are usually monocistronic and thus, promoter libraries are typically used to adjust relative expression levels ^9^. Hence, we built a set of light-responsive promoters differing in the promoter backbone and EL222 binding site number. The resulting promoters showed a wide range of maximal expression levels with promoters based on both the GAL1 and the SPO13 backbone exhibiting very low basal expression (**Fig. 2a, Supplementary Fig. 4**). However, when we analyzed the response of two promoters differing in the number of EL222 binding sites to AM, we found that they show different dose-response behaviors (**Fig. 2b**). In contrast, PWM with a period of 30 min resulted in coordinated expression with an almost linear relationship between the duty-cycle and the protein output (**Fig. 2c**). Thus, only PWM is compatible with the use of a simple promoter library for graded multi-gene expression at constant ratios (**Fig. 2d**). We observed the same behavior when both reporters were located in a single cell (**Supplementary Fig. 6a**). The use of shorter PWM periods resulted in intermediate levels of coordinated promoter regulation, allowing for input-mediated tuning of expression ratios (**Fig. 2d, Supplementary Fig. 6b**,**c for modeling results**). We note, that Elowitz and colleagues have shown that frequency modulation of TF activity can coordinate multi-gene expression in *S. cerevisiae*^10^. Thus, our work demonstrates how we can learn from natural systems to better regulate gene expression in synthetic systems using simple strategies.

**Figure 2.**
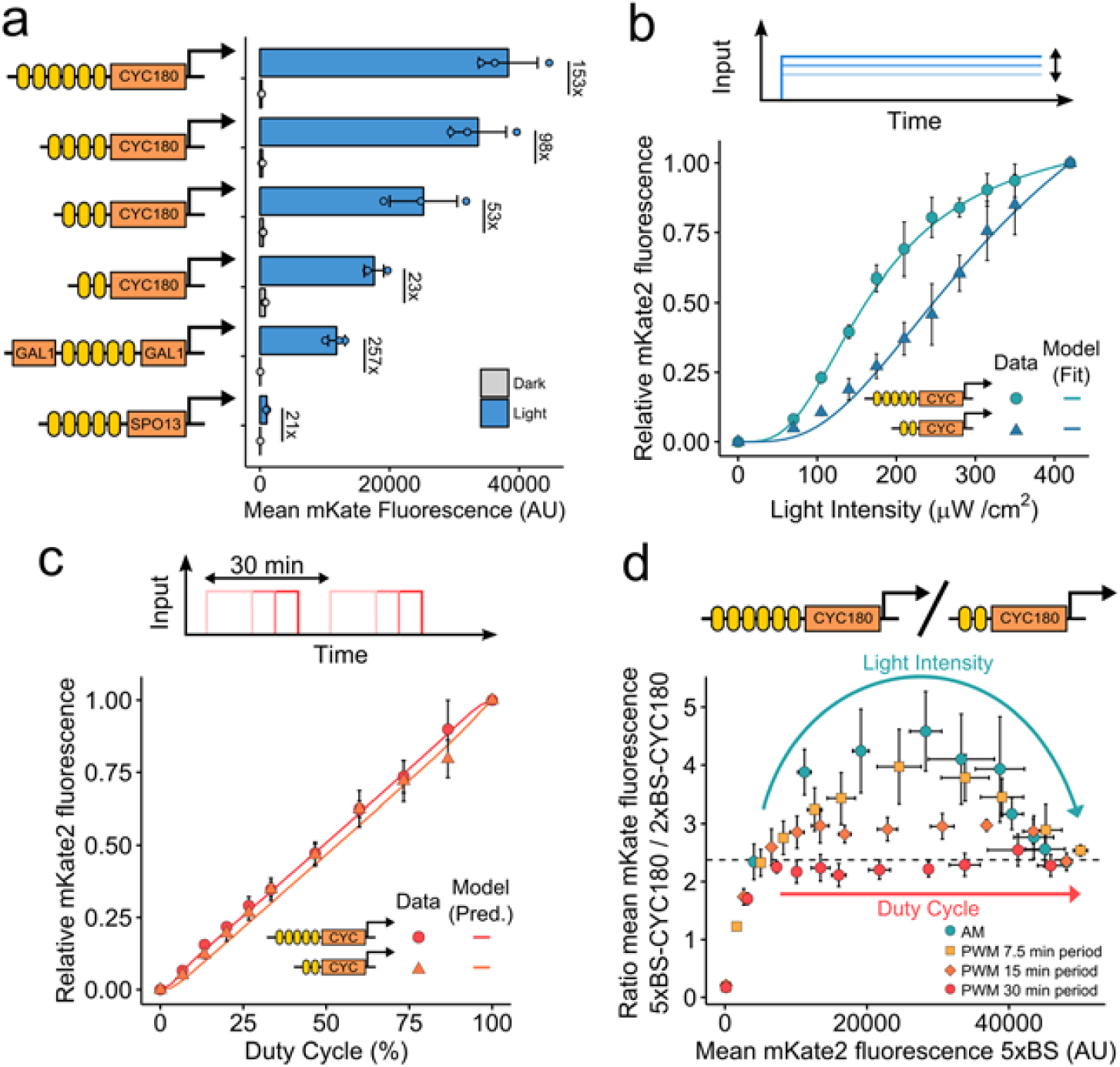
Coordinated multi-gene regulation using dynamic inputs. **(a)** A promoter library for gene expression at various expression levels. Schematics represent the different promoters. Yellow boxes represent EL222 binding sites and orange boxes represent partial sequences of yeast promoters. Strains, expressing mKate2 under the control of the respective promoter, were cultured for 6h in the dark or the presence of blue light (350 µW/cm^2^). The average cellular mKate2 fluorescence was measured using flow cytometry. Data represents the mean and s.d. of three independent experiments. **(b)** and **(c)** Dose-response of two promoters to AM (b) and PWM (c). Strains expressing mKate2 under the control of either a CYC180 promoter with five (circle, 5xBS) or two (triangle, 2xBS) VP-EL222 binding sites were grown under the illumination conditions depicted on the x-axis for 6 h. The light intensity and period for PWM were 420 µW/cm^2^ and 30 min. Mean cellular fluorescence measurements were normalized to be 0 in the dark and 1 at the highest input level to allow for easy comparison. Non-normalized values are shown in **Supplementary Fig. 6**. Data represents the mean and s.d. of three independent experiments. Lines represent model fits or predictions. (**d**) Relative gene expression levels for different induction condition. Strains (as in (b) and (c)) were grown under the same illumination conditions (light intensity and duty cycle) as shown in (b) and (c). In addition, the effect of the PWM period on coordinated expression was explored. The ratio of mKate2 expression from the 5xBS and the 2xBS promoter is plotted against the mKate2 expression from the 5xBS promoter for the same illumination conditions. The dashed line represents this ratio at constant illumination with a light intensity of 420 µW/cm^2^. Data represents the mean and s.d. of three independent experiments.

### Reducing and tuning expression variability with pulsatile signals

While we have so far only analyzed the average response of cells to input signals, gene expression can exhibit a substantial amount of heterogeneity ^11^. For many applications, precise single cell regulation of gene expression is desirable ^12^. However, the ability to tune variability may allow for the analysis of its phenotypic consequences ^11^. To date, variability regulation was achieved by the construction of synthetic gene networks ^13–16^ —namely feedback loops and cascades— as well as the tuning of promoter features, such as TATA boxes ^17^. While variability reduction via frequency modulation of TF activity was proposed theoretically, such behavior has not yet been shown experimentally ^18,19^.

For the synthetic gene expression system, PWM led to reduced cell-to-cell variability in protein levels compared to AM for the same mean expression (**Fig. 3a**). Furthermore, changing the PWM period enabled tuning of expression heterogeneity with a single input and no change in network architecture (**Fig. 3a**). In order to investigate the mechanism behind this noise reduction, we performed a dual reporter experiment (see Methods for details). This assay allows for the decomposition of expression variability stemming from stochastic events at the promoter level (intrinsic) and global differences between cells (extrinsic) ^20,21^. We found that PWM reduces both extrinsic and intrinsic variability (**Fig. 3b, c**). However, for most expression levels, extrinsic variability is the dominant source of heterogeneity in the synthetic expression system. Given that TF variability is thought to be a major determinant of extrinsic variability ^22^, we hypothesized that PWM leads to lower gene expression heterogeneity by operating in a promoter-saturating regime, where transmission of TF variability to gene expression output is minimal (**Fig. 3d**).

**Figure 3.**
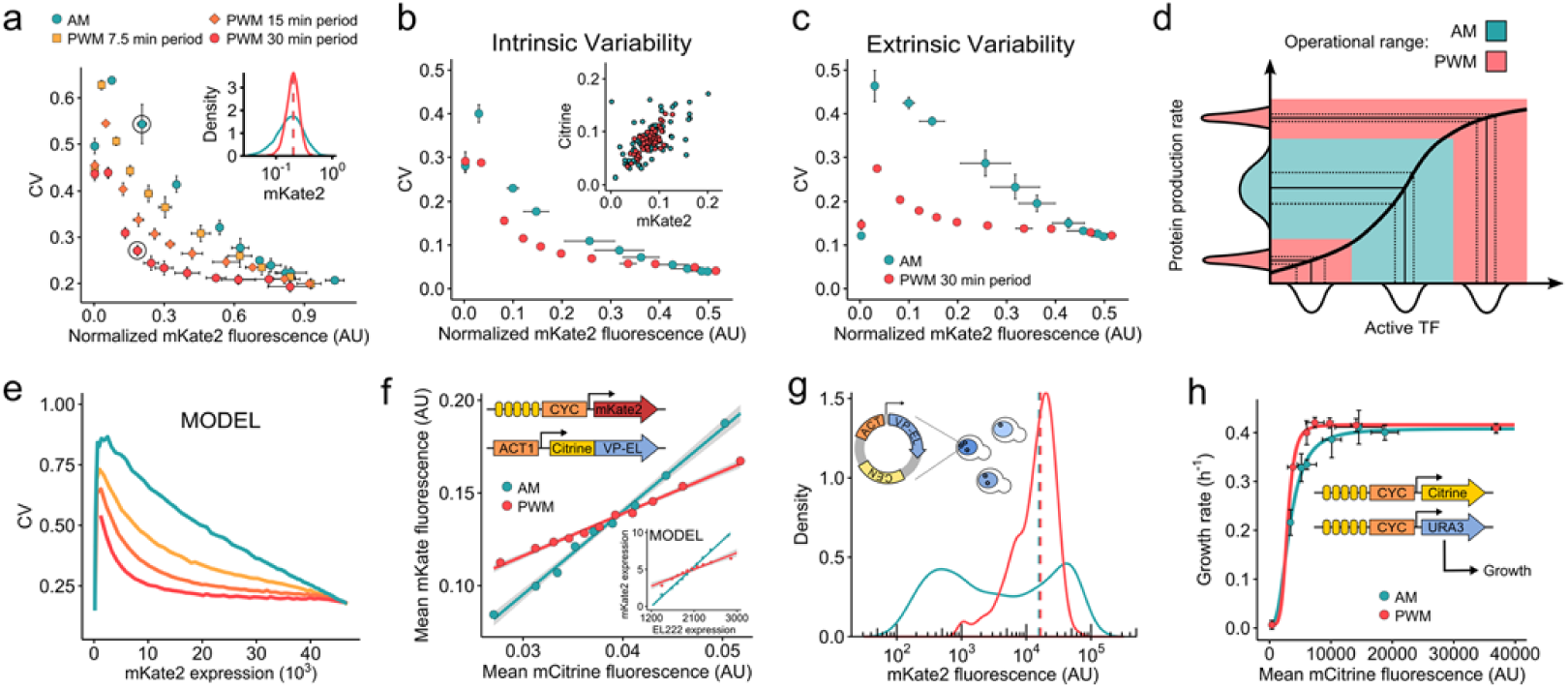
Effects of PWM and AM on gene expression variability. **(a)** Cell-to-cell variability, measured by the coefficient of variation (CV), as function of mean expression levels for different inputs. Cellular fluorescence was normalized by side-scatter measurements to reduce the influence of cell size (Methods, see **Supplementary Fig. 8** for details and non-normalized values). Cells expressing mKate2 under the control of 5xBS-CYC180pr were induced for 6 h before analysis. Illumination conditions for AM and PWM are identical to those in **Fig. 2d**. Data represents the mean and s.d. of three independent experiments. The inset shows representative fluorescence distributions for the data points circled in the main plot. **(b)** and **(c)** Contributions of intrinsic (b) and extrinsic (c) noise to gene expression variability. Experiments were performed using a diploid yeast strain based on strains expressing either the FP mKate2 or Citrine under control of 5xBS-CYC180pr from the same locus. The equivalence of both reporters is shown in **Supplementary Fig. 9**. The noise decomposition procedure is described in the Methods section. Cells were induced for 6 h under the same illumination conditions used in (a) with a 30 min PWM period. Data represents the mean and s.d. of three independent experiments. The inset shows a representative scatter plot of 60 cells for conditions with similar mean for AM and PWM (AM: 105 µW/cm^2^ light intensity, PWM: 13% duty cycle). **(d)** Schematic illustration of variability transmission from TF concentration to gene expression. The graph represents the input-output function of active TF concentration to protein production rate (black line). Histograms represent cell-to-cell variability of both species. Assuming that the input-output function is identical between cells, cell-to-cell variability in gene expression rate increases with the steepness of the input-output function. In contrast to AM, PWM operates mainly at the extremes of this relationship where the slope is minimal, effectively leading to reduced variability transmission. **(e)** CV as function of mean expression levels as calculated using the 5xBS-CYC180pr model including VP-EL222 variability is shown for different inputs. Induction regimes are as in (a). See **Supplementary Note 1.6** for discussion and details of the modeling and **Supplementary Fig. 10b**,**c** for model predictions and data for 2xBS-CYC180pr. (**f**) Dependence of gene-expression output on mCitrine-VP-EL222 levels for AM and PWM. Fluorescence was analyzed after 2 h of induction to retain a causal relationship between instantaneous mCitrine-VP-EL222 and mKate2 levels (CV-mean relationship is shown in **Supplementary Fig. 10d**). Cells were collected in 10 bins of equal cell number based on their normalized mCitrine fluorescence. Data points represents the mean normalized mKate2 and mCitrine fluorescence of cells from each bin. Lines represent results of linear regressions. The inset shows the model prediction for conditions leading to similar relative gene expression levels. (**g**) Effect of AM (blue) and PWM (red) on fluorescence distributions with VP-EL222 expressed from a centromeric plasmid. Induction conditions: AM = 105 µW/cm^2^ light intensity, PWM = 420 µW/cm^2^ light intensity; 45 min period; 26.7 % duty cycle. CV-mean relationship for other conditions is shown in **Supplementary Fig. 11**. (**h**) Effect of regulating URA3 expression levels by AM and PWM on cell growth. Cells expressing both mCitrine and URA3 from 5xBS-CYC180pr were grown under light induction for 13 h in media with uracil before transfer to uracil-free medium, fluorescence, and growth measurements. Hill functions were fit to the data for guidance (see **Supplementary Table 5** for parameters). Effect of AM and PWM on the CV is shown in **Supplementary Fig. 12**. Data represents the mean and s.d. of two independent experiments.

To approximate this phenomenon with our simple mathematical model, we performed simulations in which we drew TF concentration from a log-normal distribution describing the single-cell distribution of a mCitrine-tagged ^23^ version of VP-EL222 (**Supplementary Note 1.6, Supplementary Fig. 10a**). This model can qualitatively recapitulate the experimental data (**Fig. 3e, Supplementary Fig. 10b**,**c**). We further showed experimentally that PWM reduced the slope of the correlation between VP-EL222 expression levels and mKate2 output (**Fig. 3f, Supplementary Fig. 10d**). Next, we expressed VP-EL222 from a centromeric plasmid to increase TF variability by introducing plasmid copy number variation (**Supplementary Fig. 11**) ^24^. Under these conditions, AM led to a wide-spread multi-modal protein distribution at intermediate expression levels (**Fig. 3g**). In contrast, PWM resulted in the merging of these distributions. Thus, the use of PWM does not only reduce heterogeneity as measured by the CV but may also lead to qualitatively different distributions by attenuating the effects of TF variability on downstream gene expression.

Finally, we sought to map the expression level of the metabolic enzyme URA3 on cell growth in the absence of uracil. We found that the dose response of mean expression to growth depends on expression heterogeneity with tight regulation enabling maximal growth at lower expression levels (**Fig. 3h**). This result exemplifies the importance of precise regulation for the analysis of expression-phenotype relationships and for the adjustment of optimal protein levels for synthetic biology applications in which metabolic burden is non-negligible ^25^.

## DISCUSSION

We present a highly inducible, fast-acting optogenetic expression system for *S. cerevisiae* which enables the regulation of protein levels by PWM. Learning from the use of pulsatile regulation in a natural system ^10^, we show that PWM enables the use of simple promoter libraries and a single input for the graded and coordinated regulation of multiple genes at different expression levels. We further uncover a novel mechanism for noise reduction and tuning in gene expression systems based on pulsatile inputs. Thus, the simple use of dynamic inputs may replace laborious optimization of promoter dose-response curves ^26^ and construction of gene networks ^14,17^ for a variety of synthetic biology applications. Future work may show whether pulsatile regulation is employed for the control of cell-to-cell variability in natural systems. Notably, the mechanisms behind the benefits of pulsatile regulation are not specific to VP-EL222 and should be widely applicable to systems based on fast-acting regulators in a variety of organisms. For example, attenuation of TF variability may be important for the precise and graded control of endogenous transcription using light-inducible CRISPR-Cas9 systems ^27^ in mammalian cells, where transient transfections are often performed.

## Acknowledgments

We thank Peter Buchmann and Marc Rullan for the construction of and help with the LED-setup. We thank members of the Khammash lab, Serge Pelet, and Martin Ackermann for helpful discussion and Stephanie Aoki for critical reading of the manuscript. We would further like to thank Fabian Rudolf (ETH Zurich), Kevin Gardner (City University of New York), Robert H Singer (Albert Einstein College of Medicine), and Daniel Zenklusen (Université de Montréal) for providing plasmids. DB is part of the Life Science Zurich graduate school.

## AUTHOR CONTRIBUTIONS

D.B. conceived the study, performed experiments and mathematical modeling, analyzed data, and wrote the paper. M.K. supervised the study, analyzed data, and wrote the paper.

## Data and code availability statement

Data, plasmids, strains, and custom code are available from the corresponding author upon request.

## Materials and Methods

### Plasmid construction

*E. coli* TOP10 cells (Invitrogen) were used for plasmid cloning and propagation. Sequences and details of all DNA constructs used in this study can be found in **Supplementary Note 2**. All plasmids used in this study are summarized in **Supplementary Table 1**. Plasmids were constructed by restriction-ligation cloning using enzymes from New England Biolabs (USA). All PCRs were performed using Phusion Polymerase (New England Biolabs). The EL222-based transcription factor under control of the ACT1 promoter was cloned in an integrative vector based on the pRS vector series and a low-copy plasmid (pRG215) ^28^. Constructs with light inducible promoters were cloned in pFA6a-His3MX6 ^29^. All constructs were verified by sanger sequencing (Microsynth AG, Switzerland).

### Yeast strain construction

All strains are derived from BY4741 and BY4742 (Euroscarf, Germany) ^30^. All strains used in this study are summarized in **Supplementary Table 2**. Transformations were performed with the standard lithium acetate method ^31^ and selection was performed on appropriate selection plates. The basis of the majority of strains used in this study are DBY41 and DBY42. Both strains express VP-EL222 from the ACT1 promoter and were generated by transforming PacI digested plasmid pDB58 into BY4741 and BY4742 respectively. Plasmid integration was verified by function. All light inducible promoter constructs were PCR amplified using primers for the integration into the HIS3 locus (Primers HIS3_integration_fwd/rv, **Supplementary Table 3**). Integration of reporter constructs was verified via PCR and function. Diploid strains were generated by mating and selection by growth on SD plates lacking both L-Lysine and L-Methionine.

### Media and growth conditions

All experiments were performed in synthetic medium (SD; LOFLO yeast nitrogen (ForMedium), 5 g/L ammonium sulfate, 2% glucose, pH was adjusted to 6.0). All experiments were performed in 25 ml glass centrifuge tubes (Schott 2160114, Duran) stirred with 3 × 8 mm magnetic stir bars (Huberlab) using a setup comprised of a water bath (ED (v.2) THERM60, Julabo) set to 30 °C, a multi position magnetic stirrer (Telesystem 15, Thermo Scientific) set to 900 rpm, a 3D printed, custom-made 15-tube holder, and custom-made LED pads located underneath the culture tubes. A white diffusion filter (LEE Filters) was placed between the LED and the culture tube to allow for even illumination. LED intensity was measured at 4 cm distance from the light source using a NOVA power meter and a PD300 photodiode sensor (Ophir Optronics).

### Flow cytometry

All experiments except growth and smFISH were performed in the following way.

Cultures were grown overnight starting from single yeast colonies, subcultured in fresh medium and grown for at least 16 h in the dark while maintaining an optical density at 700 nm (OD_700_) lower than 0.4. At the start of the experiment, cells were diluted to an OD_700_ of 0.005 in 4 ml of medium. Before measurement, cell samples were incubated in SD with 0.1 mg/ml cycloheximide for 3.5 h at 30 °C to ensure full fluorescent protein maturation. Samples were analyzed using a LSRFortessaTM LSRII cell analyzer (BD Biosciences, Germany). To measure mKate2 fluorescence, a 561 nm excitation laser and a 610/20 nm emission filter and for mCitrine, a 488 nm excitation laser and a 530/11 nm emission filter were used. Data was analyzed using R with the flowCore package ^32^. Cells were gated based on forward and side scatter to remove debris and cell aggregates. For strains containing centromeric plasmids, a budded cell population was selected by gating based on the forward and side scatter width ^33^. We found that this population shows a higher percentage of responsive cells, which likely results from a higher degree of plasmid retention. Strong outliers were removed from the data as follows: First, the fluorescence values were log-transformed. Outliers were defined as data-points with an absolute deviation from the fluorescence distribution median of greater than 3-fold of the median absolute deviation.

For the analysis of gene expression heterogeneity, fluorescent levels were normalized by side scatter area to reduce the effect of cell size (see **Supplementary Fig. 8**) ^34^. At least 1000 cells per sample were analyzed.

### Modeling

The modeling and parameter fitting is described in detail in **Supplementary Note 1**. The model consists of the following three ordinary differential equations describing VP-EL222 activation (1), VP-EL222 dependent mRNA expression (2), and protein expression (3):

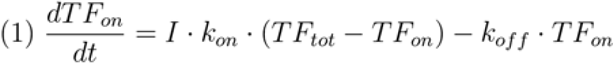

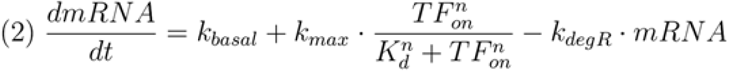

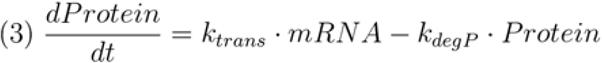

Simulations and model fitting were performed using Matlab (R2014a, Mathworks).

### Dual-reporter experiments

Dual-reporter experiments were performed using the diploid strain DBY110. This strain was constructed by mating DBY43 and DBY104, expressing mKate2 and mCitrine from 5xBS-CYC180pr integrated into the HIS3 locus. The equivalence of both reporter genes is shown in **Supplementary Fig. 9**. mCitrine fluorescence values were adjusted by multiplication with a constant in order to equate the mean values of mCitrine and mKate2 fluorescence measurements. Using the formalism introduced in Ref. ^35^, total variability was decomposed into extrinsic and intrinsic variability using the following equations:

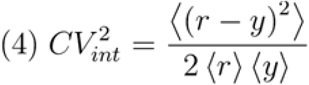

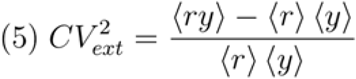

Here, r and y are vectors whose elements are cellular fluorescence values for mKate2 and mCitrine, respectively. Angled brackets represent population means.

### Measuring the influence of light on cell growth

Cells were initially grown as described for flow cytometry experiments. At the start of the experiment, cultures were diluted to an OD_700_ of 0.01 in a total volume of 6 ml. Cells were grown for 2 h before starting blue-light illumination. Subsequently the OD_700_ was measured every hour for 6 h and the growth rate was calculated by performing linear regressions on log-transformed OD-data.

### Measuring the effect of Ura3 on cell growth

In order to measure how Ura3 expression affects cell growth, DBY125 expressing both mCitrine and Ura3 from two separate 5xBS-CYC180 promoters was initially grown as described for all other flow cytometry experiments (see above). Cells were then diluted to an OD of 0.001 in SD medium and illuminated for 13 hours under the following light conditions. AM: 0, 63, 70, 77, 84, 98, 126 µW/cm^2^ light intensity. PWM: 420 µW/cm^2^ light intensity; 30 min period; 5, 6.6, 8.3, 10, 16.6, 50 % duty cycle.

Cells were then washed two times with SD medium lacking L-uracil and were diluted 1:10 in 5 ml of this medium. Cells were further grown under the same illumination conditions and samples were analyzed every hour by measuring the cell count per 57 µl medium using an Accuri C6 flow cytometer (BD Biosciences) for 7 h. Growth rates were calculated by performing linear regressions on log-transformed count-data. Cellular mCitrine fluorescence was measured as described above after the transfer to uracil-free medium. The use of mCitrine expression from an independent promoter as a proxy for Ura3 expression is permitted by the fact that URA3 is a stable protein ^36^.

### Single molecule FISH experiments

For single molecule FISH experiments, DBY89 was grown from a single colony to saturation in SD medium. Cultures were diluted to reach an OD_700_ of 0.4 at the start of the experiment the next day. For each time point, 4 ml cell culture were transferred to 25 ml glass centrifuge tubes stirred with 3 × 8 mm magnetic stir bars. Illumination was performed with a light intensity of 350 µW/cm^2^ for 20 min.

Cell fixation and probe hybridization was performed as described previously ^37^. Briefly, after 0, 10, 20, 30, 40, and 60 min (where 0 min marks the start of illumination), cells were fixed for 45 min after adding 400 µl of 37% formaldehyde (Sigma Aldrich) to the culture medium. Spheroplasting was performed using a final Lyticase (Sigma-Aldrich) concentration of 50 Units/ml. The progression of spheroplasting was monitored under the microscope. Cells were stored in 70% ethanol at 4 °C overnight. Hybridization was performed using multiple probes complementary to the PP7 SL and singly labeled with CY3 at a 0.1 µM concentration (synthesized by Integrated DNA Technologies, sequences are listed in **Supplementary Table 3**).^38^ Cells were stained with DAPI (0.1 μg/ml in PBS, Sigma-Aldrich), attached to Poly-D-Lysine treated coverslips, and slips were mounted on slides using Prolong Gold mounting medium (Invitrogen).

### Microscopy Setup

All images were taken with a Nikon Ti-Eclipse inverted microscope (Nikon Instruments), equipped with a Plan Apo Lambda 100X Oil objective (Nikon Instruments), Spectra X Light Engine fluorescence excitation light source (Lumencor, USA), pE-100 brightfield light source (CoolLED Ltd., UK), and CMOS camera ORCA-Flash4.0 (Hamamatsu Photonic, Switzerland). The camera was water-cooled with a refrigerated bath circulator (A25 Refrigerated Circulator, Thermo Scientific). The microscope was operated using NIS-Elements software. Z-stacks consisting of 31 images with a step size of 0.1 µm were taken for CY3 (Excitation: 542/33, Emission: 595/50) and DAPI (Excitation: 390/22, Emission: 460/50). Phase contrast images were taken at the reference point.

### Microscopy image analysis

The image analysis procedure was performed using custom Matlab scripts and consists of three steps: segmenting individual nuclei (based on DAPI images), locating fluorescent spots in the nuclear regions, and quantifying the intensity of these spots.

Nuclei were first enhanced by using the difference of Gaussians algorithm. Nuclear regions were then segmented by manually optimized thresholding. Detected regions that were too big or small to represent nuclei were removed. For each nuclear region, a Difference of Gaussian algorithm was used to enhance spots in the CY3 images and spots were identified using thresholding. In order to quantify the intensity of the nuclear spots, the sum of a two-dimensional Gaussian function and a 2D-plane was fitted in a square area around the identified spot with an edge length of 19 pixels. If no spot was detected, the same procedure was performed at the center of the nuclear region. Spot intensity was then defined as the integral of the Gaussian function. For each nucleus / cell, the spot with the highest intensity was defined as the transcription site.

## Supplementary Information

### Supplementary Figures

**Supplementary Figure 1.**
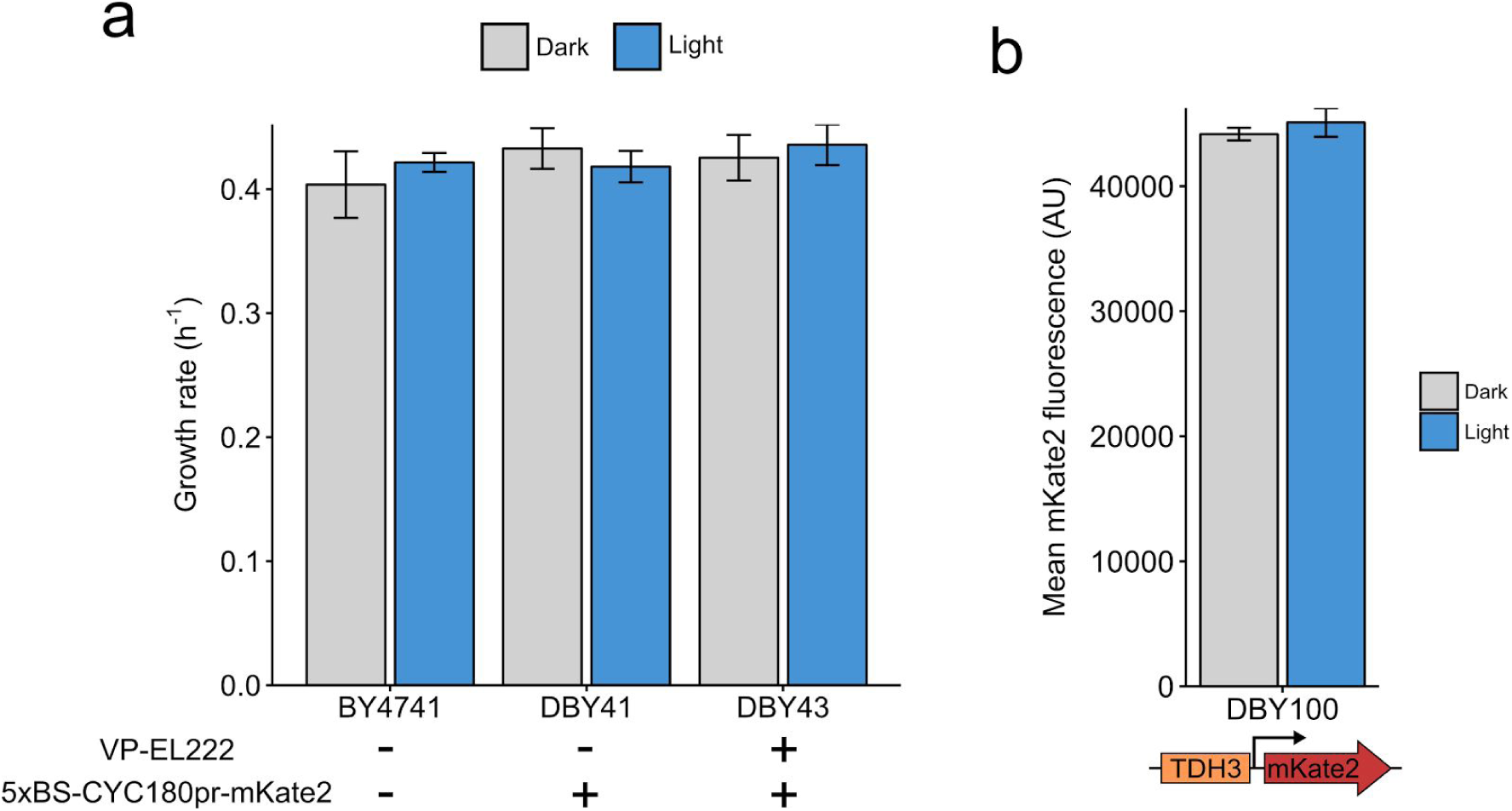
VP-EL222 expression and illumination does not affect cell growth and constitutive gene expression. (**a**) Yeast strains, with or without the VP-EL222 and a reporter construct (5xBS-CYC180pr-mKate2), were grown in the dark or under blue light illumination (420 µW/cm^2^) for 6 h. The OD_700_ was measured every hour and the growth rate was calculated. Data represents the mean and s.d. of two independent experiments. The average growth rate of all experiments is 0.42 h^-1^. This value was used as the protein dilution rate in the mathematical modeling. (**b**) A yeast strain expressing mKate2 constitutively from the TDH3 promoter (DBY100) was grown either in the dark or under illumination (420 µW/cm^2^) for 6 h. The average cellular mKate2 fluorescence was measured using flow cytometry. Data represents the mean and s.d. of two independent experiments.

**Supplementary Figure 2.**
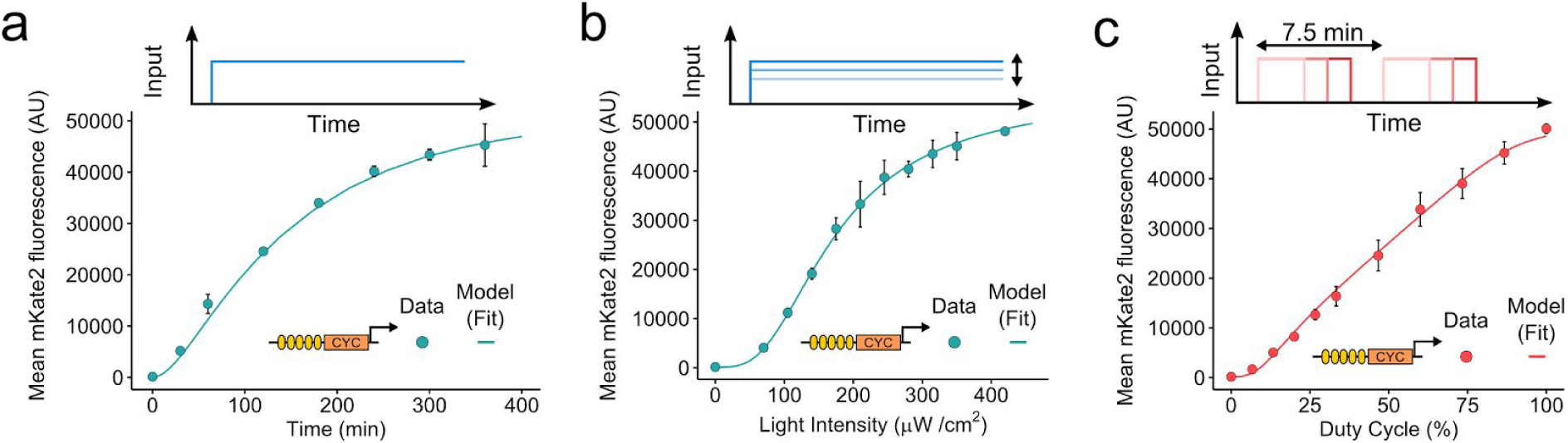
Characterization experiments and results of model fitting. Three characterization experiments were performed with DBY43 (containing the VP-EL222 and a reporter construct (5xBS-CYC180pr-mKate2)). Data points represent mean cellular mKate2 fluorescence (measured by flow cytometry, mean and s.d. of three independent experiments) and lines represent the model fits. (**a**) Time-course of VP-EL222 mediated gene expression. Cells were illuminated with blue light (350 µW/cm^2^) for 6 h and fluorescence was measured at regular intervals. (**b**) Dose response to AM / light intensity. Fluorescence was measured after 6 h of illumination. (**c**) Dose response to PWM with a 7.5 min period / duty cycle. The light intensity was 420 µW/cm^2^.

**Supplementary Figure 3.**
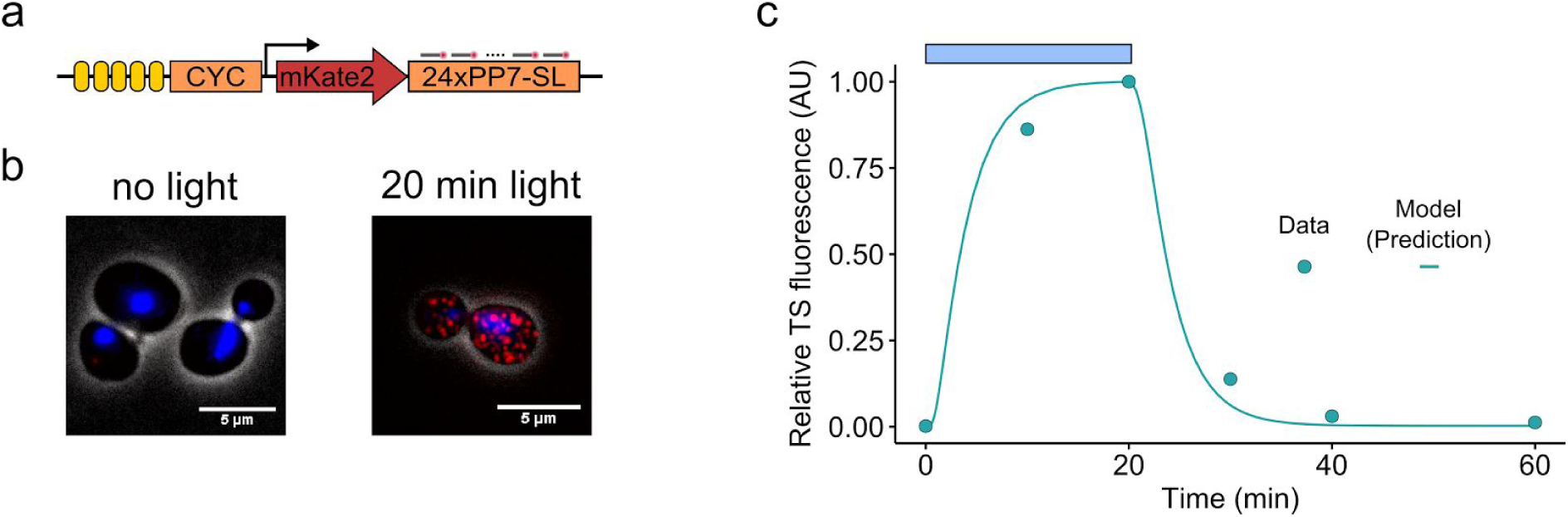
smFISH analysis of VP-EL222 mediated transcription. A yeast strain containing VP-EL222 and a reporter construct containing a sequence coding for 24 copies of the PP7 stem loop (5xBS-CYC180pr-mKate2-24xPP7-SL) were grown under blue light illumination (420 µW/cm^2^) for 20 min and subsequently in the dark for 40 min. Samples were taken before illumination and after 10, 20, 30, 40, 60 min. smFISH was performed with CY3 labeled probes complementary to the PP7-SL (probe sequences are listed in **Supplementary Table 3**). (**a**) Schematic of the reporter construct. (**b**) Representative microscopy images before and after 20 min of illumination. Grayscale: phase contrast / cell boundaries, blue: DAPI channel (maximum intensity projection), red: CY3 channel (maximum intensity projection). (**c**) Time-course of nascent RNA quantification. Points represent measured fluorescence at the transcription site (TS) relative to the maximal value at 20 min (see Methods for details on the quantification). The line represents the model prediction for nascent RNA accumulation (see **Supplementary Note 1.4** for modeling details).

**Supplementary Figure 4.**
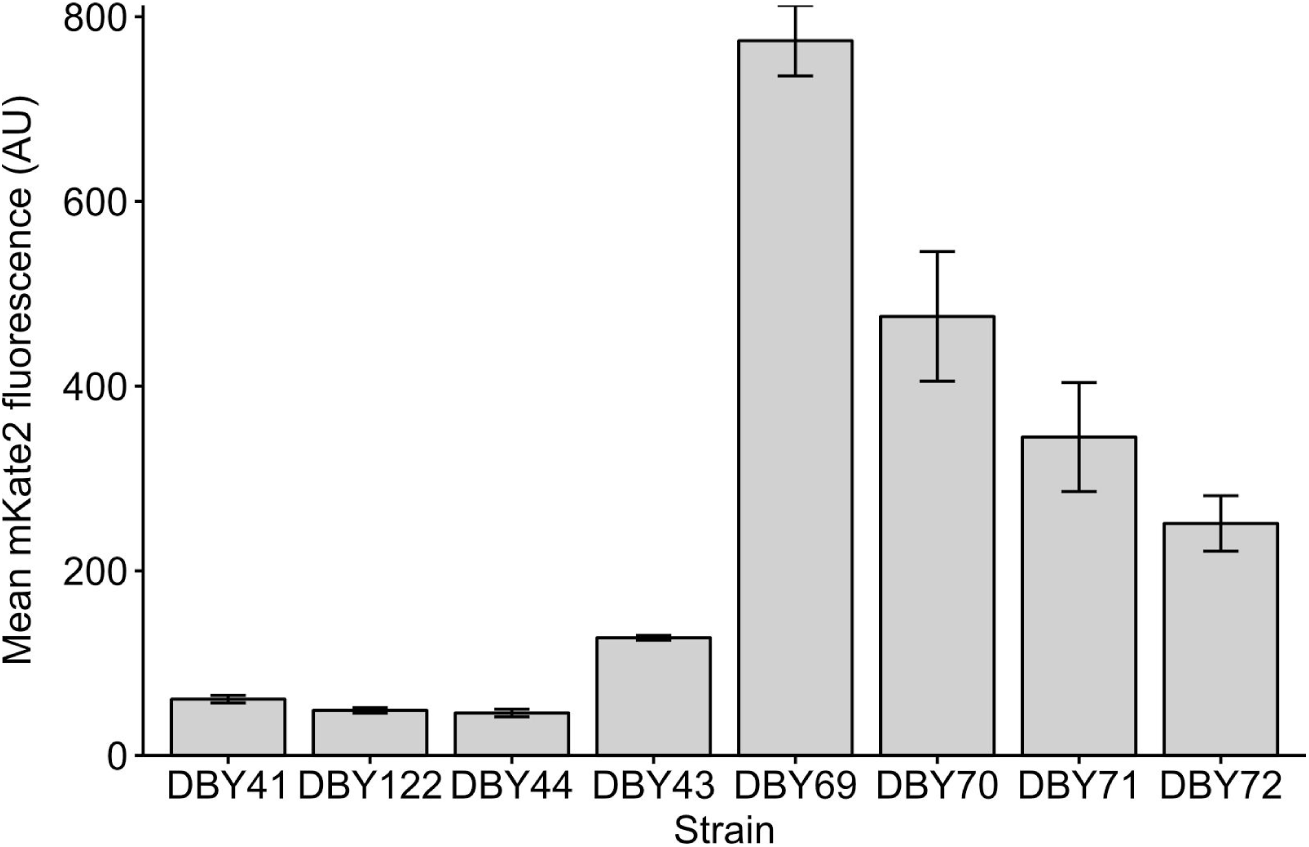
Comparison of expression from different VP-EL222 regulated promoters in the dark. Strains, expressing mKate2 under the control of different promoters (see **Supplementary Table 2**), were cultured for 6h in the dark (350 µW/cm^2^). The average cellular mKate2 fluorescence was measured using flow cytometry. Data represents the mean and s.d. of three independent experiments.

**Supplementary Figure 5.**
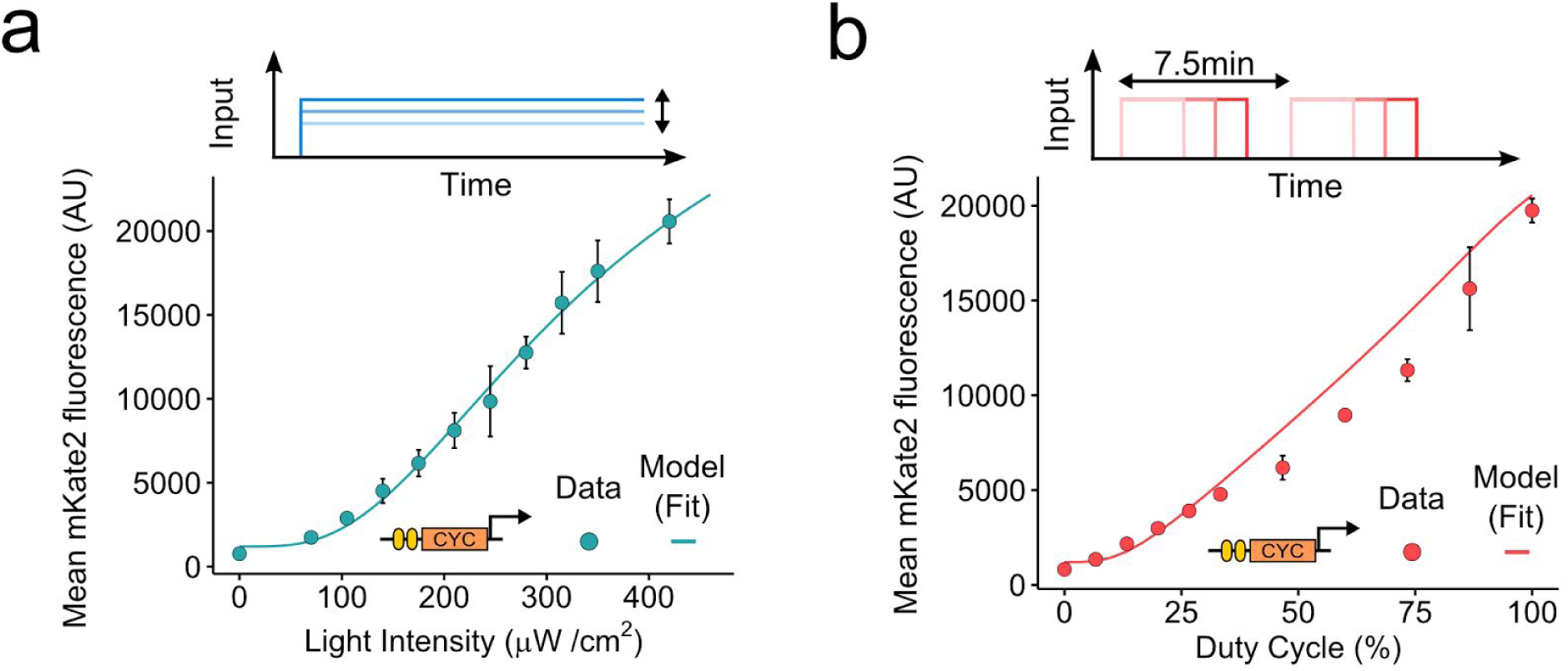
Model fits for a VP-EL222 regulated promoter based on the CYC1 promoter with two VP-EL222 binding sites (2xBS-CYC180pr, DBY69). Data points represent mean cellular mKate2 fluorescence (measured by flow cytometry after 6 h of illumination, mean and s.d. of three independent experiments) and lines represent the model fits. (**b**) Dose response to AM / light intensity. (**c**) Dose response to PWM / duty cycle. The PWM period was 7.5 min. The light intensity was 420 µW/cm^2^.

**Supplementary Fig. 6.**
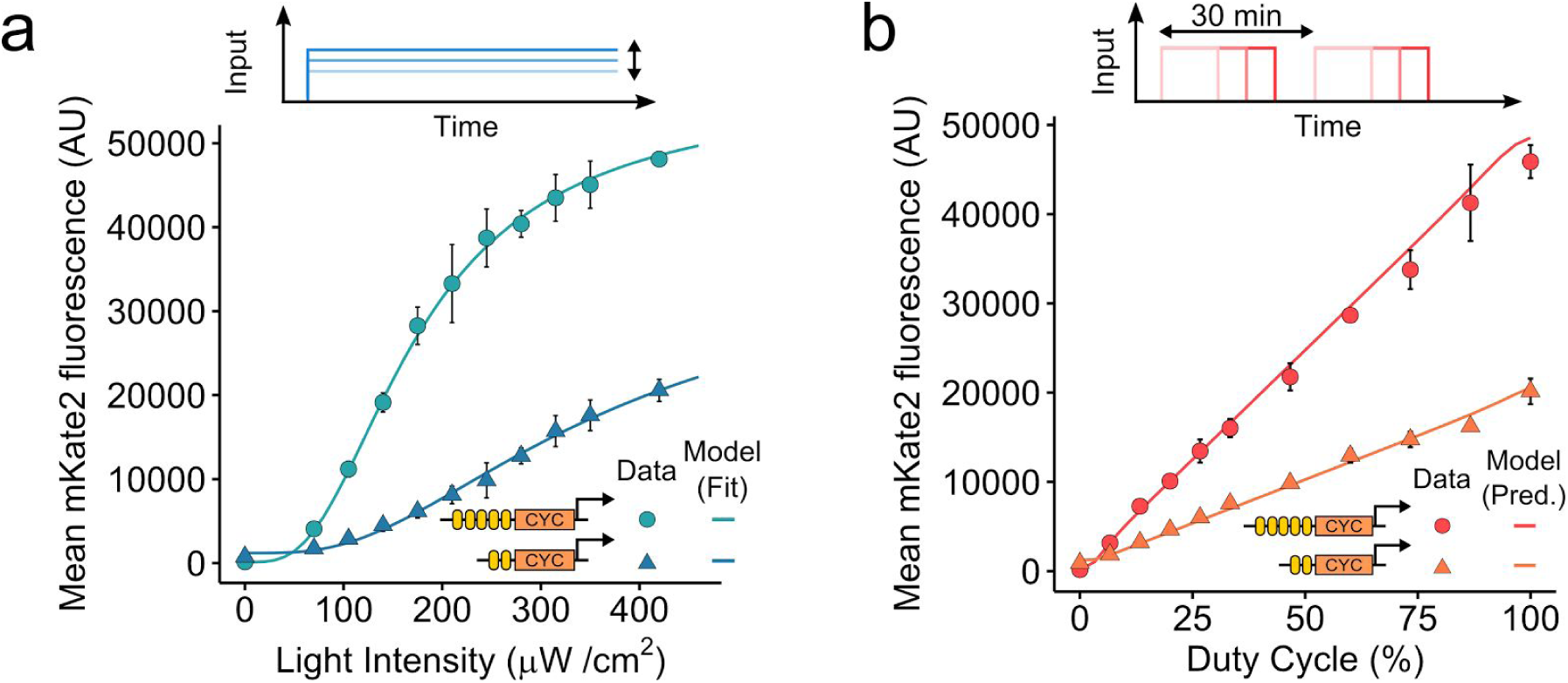
Non-normalized curves showing the coregulation of 2xBS-CYC180pr and 5xBS-CYC180pr. Dose-response of the two promoters to AM (**a**) and PWM (**b**). Strains expressing mKate2 under the control of either a CYC180 promoter with five (circle, 5xBS) or two (triangle, 2xBS) VP-EL222 binding sites were grown under the illumination conditions depicted on the x-axis for 6 h. The light intensity and period for PWM were 420 µW/cm^2^ and 30 min. Data represents the mean and s.d. of three independent experiments. Lines represent model fits or predictions.

**Supplementary Fig. 7.**
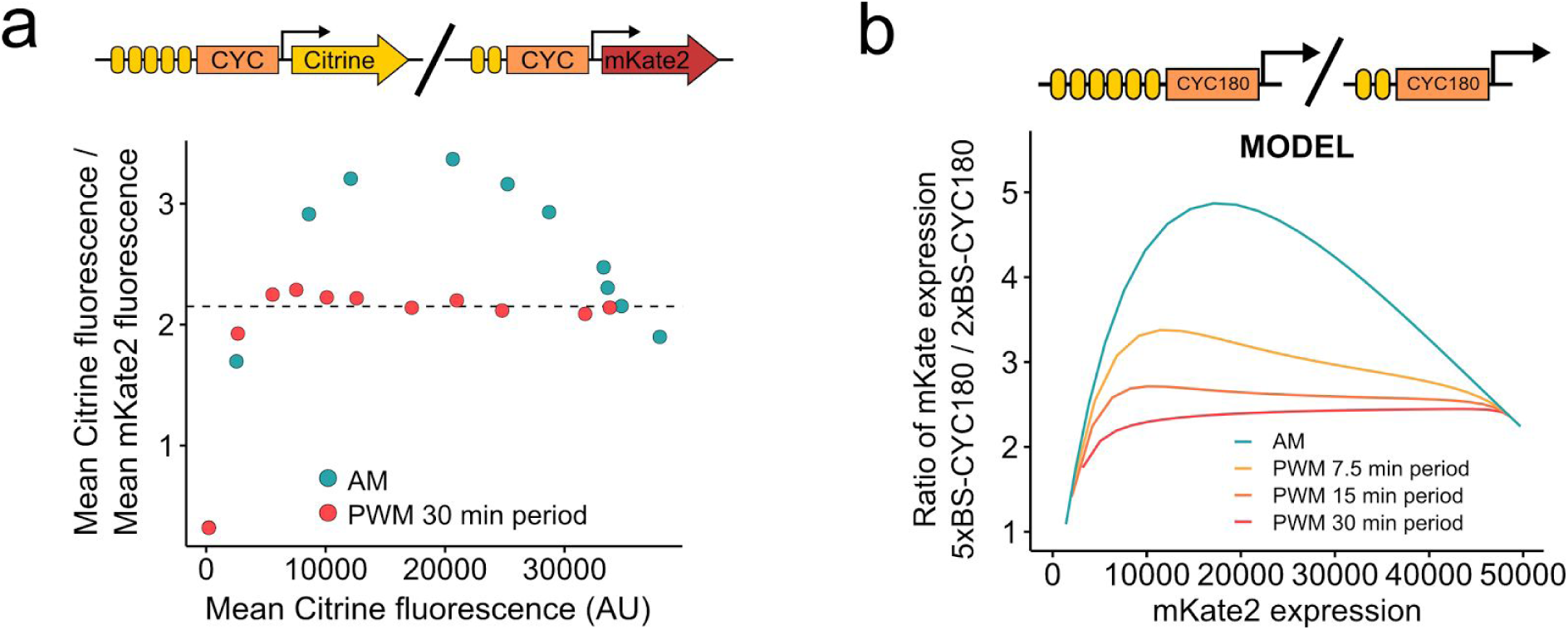
Further results on PWM mediated co-regulation. (**a**) Co-regulation of 5xBS-CYC180pr-mCitrine and 2xBS-CYC180pr-mKate2 in single cells. The diploid yeast strain DBY118 was grown under different induction conditions for 6h and fluorescence was analyzed using flow-cytometry. PWM was performed with a 30 min period and a light intensity of 420 µW/cm^2^. AM was performed with the following light intensities: 105, 140, 175, 210, 245, 280, 315, 350, 420, 490 µW/cm^2^. The dashed line represents this ratio at constant illumination with a light intensity of 420 µW/cm^2^. (**b**) Model results for effect of AM and PWM with different periods on co-regulation of gene expression from 5xBS-CYC180pr and 2xBS-CYC180pr.

**Supplementary Fig. 8.**
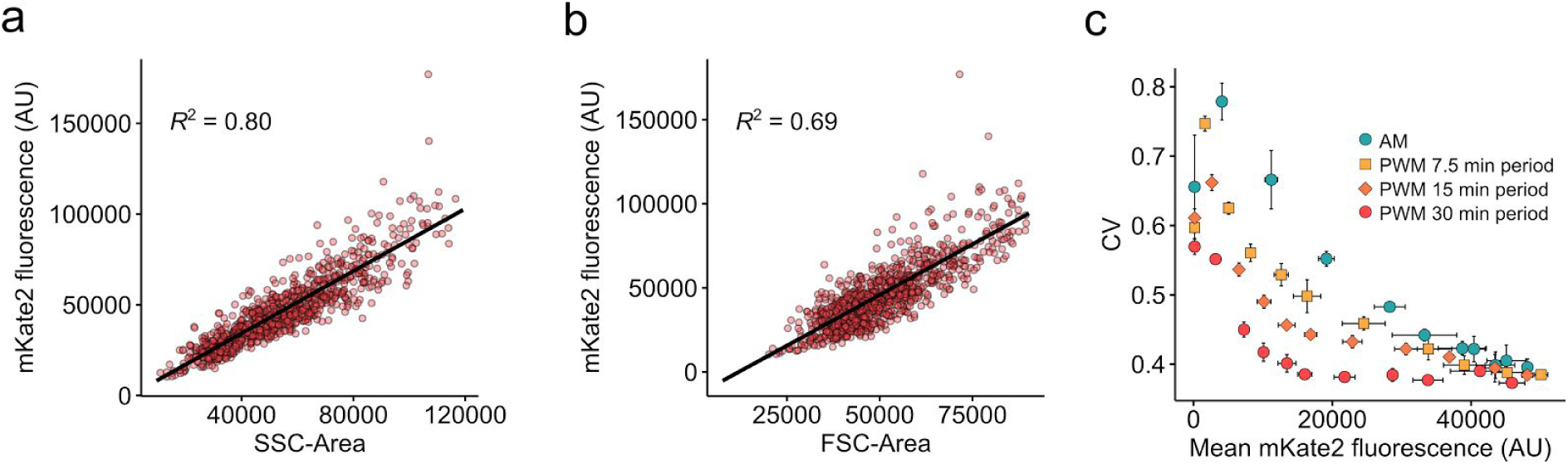
Normalization of cellular fluorescence by scatter as a proxy for cell size. Differences in cell size can lead to strong cell-to-cell variability in cellular protein abundances that can mask the magnitude of other variability sources. We thus analyzed the correlation of side scatter (**a**, SSC) and forward scatter (**b**, FSC) with constitutive mKate2 expression from the TDH3 promoter. Points represent individual cells. Lines represent linear regression with the R^2^ values shown on the graph. We found that both SSC and FSC correlate linearly with gene expression. Due to the stronger correlation with SSC, we decided to normalize fluorescence values by dividing, for each cell the fluorescence readout by the SSC readout. (**c**) CV as function of mean expression levels from 5xBS-CYC180pr for different inputs. The experiment is the same as shown in **Fig 3a**, without normalization by SSC. The data shows that the qualitative relationships are not affected by normalization.

**Supplementary Figure 9.**
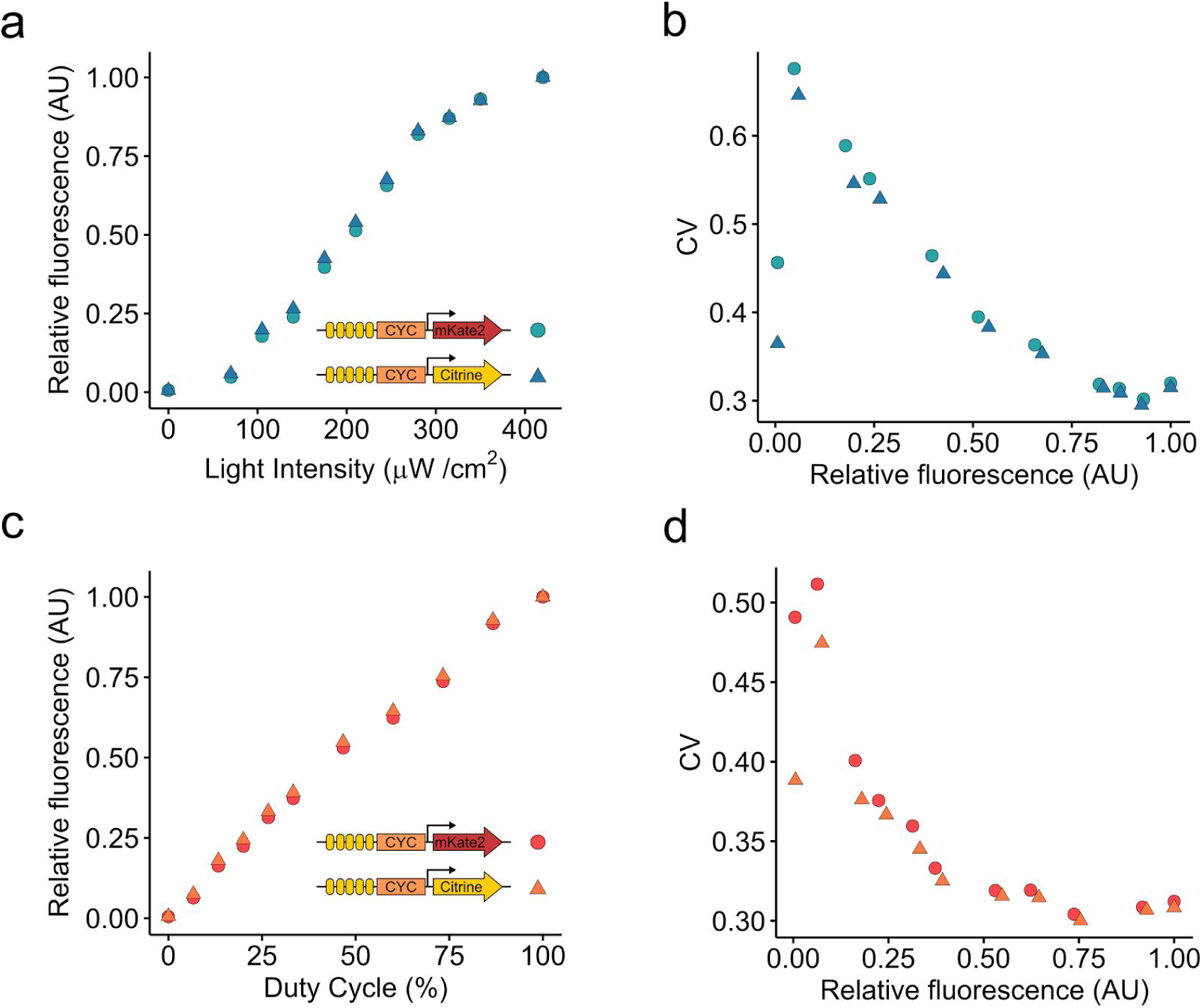
Equivalence of mKate2 and mCitrine reporters used for dual-color noise decomposition. AM (**a**,**b**) and PWM (**c**,**d**) experiments were performed with the diploid yeast strain DBY110 expressing both mKate2 and mCitrine from 5xBS-CYC180pr. Dose response curves (**a**,**c**) were normalized by dividing each measured value by the maximum value obtained for a given FP in order to reach equivalent fluorescence values for mKate2 and mCitrine. Cells were illuminated under the conditions depicted on the x-axis for 6 h. The light intensity and period for PWM were 420 µW/cm^2^ and 30 min. (**b**,**d**) CV is plotted against mean expression for the same experiment shown in (a) and (c) to analyze the equivalence of FP distributions. The CV differs under non-induced conditions. This is likely results from differences in cellular autofluorescence in both fluorescence channels.

**Supplementary Fig. 10.**
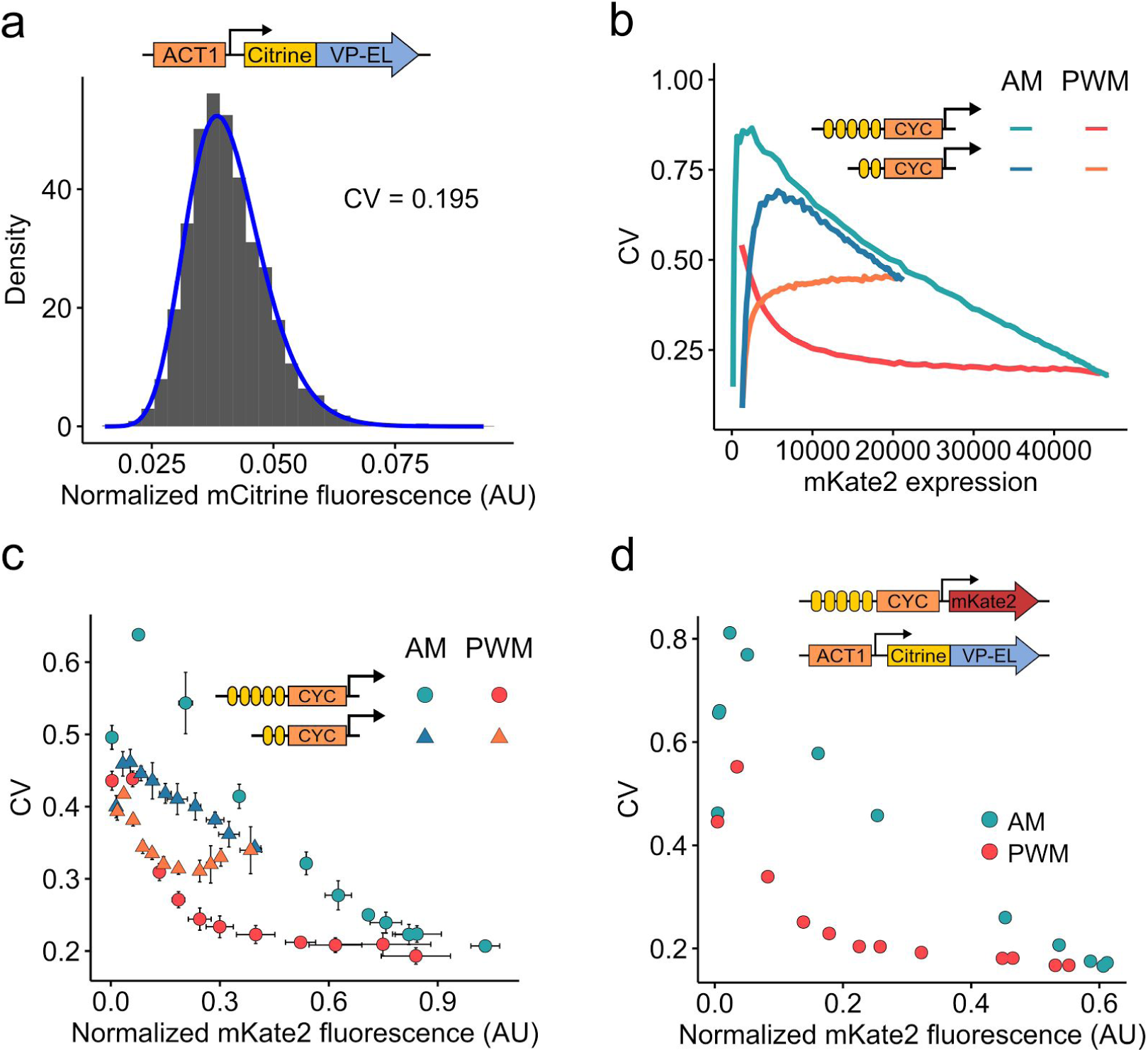
Measurement and modeling of VP-EL222 variability. (**a**) Distribution of SSC-normalized mCitrine fluorescence as measured by flow cytometry. Cells express mCitrine-VP-EL222 under control of the ACT1 promoter. The line represents the fit of a lognormal distribution to the data. The CV of the fitted distribution is shown on the graph. (**b**) The CV-mean expression relationships as calculated from the model for 5xBS-CYC180-pr and 2xBS-CYC180pr are shown. The response to AM (blue) and PWM (red/orange) was evaluated. PWM conditions: 30 min period, 420 µW/cm^2^ light intensity. (**c**) The CV is plotted as a function of mean expression from 5xBS-CYC180pr (circle) and 2xBS-CYC180pr (triangle) in response to AM (blue) and PWM (red/orange). For 5xBS-CYC180pr, results and induction conditions are the same as in **Fig. 3a** with a 30 min PWM period. Results for 2xBS-CYC180pr were obtained under the same conditions. Data represents the mean and s.d. of three independent experiments. (**d**) The CV is plotted as a function of mean expression for mCitrine-VP-EL222 mediated expression from 5xBS-CYC180-pr under AM and PWM. Cells were analyzed after 2h of induction. The light intensity and period for PWM were 420 µW/cm^2^ and 30 min.

**Supplementary Fig. 11.**
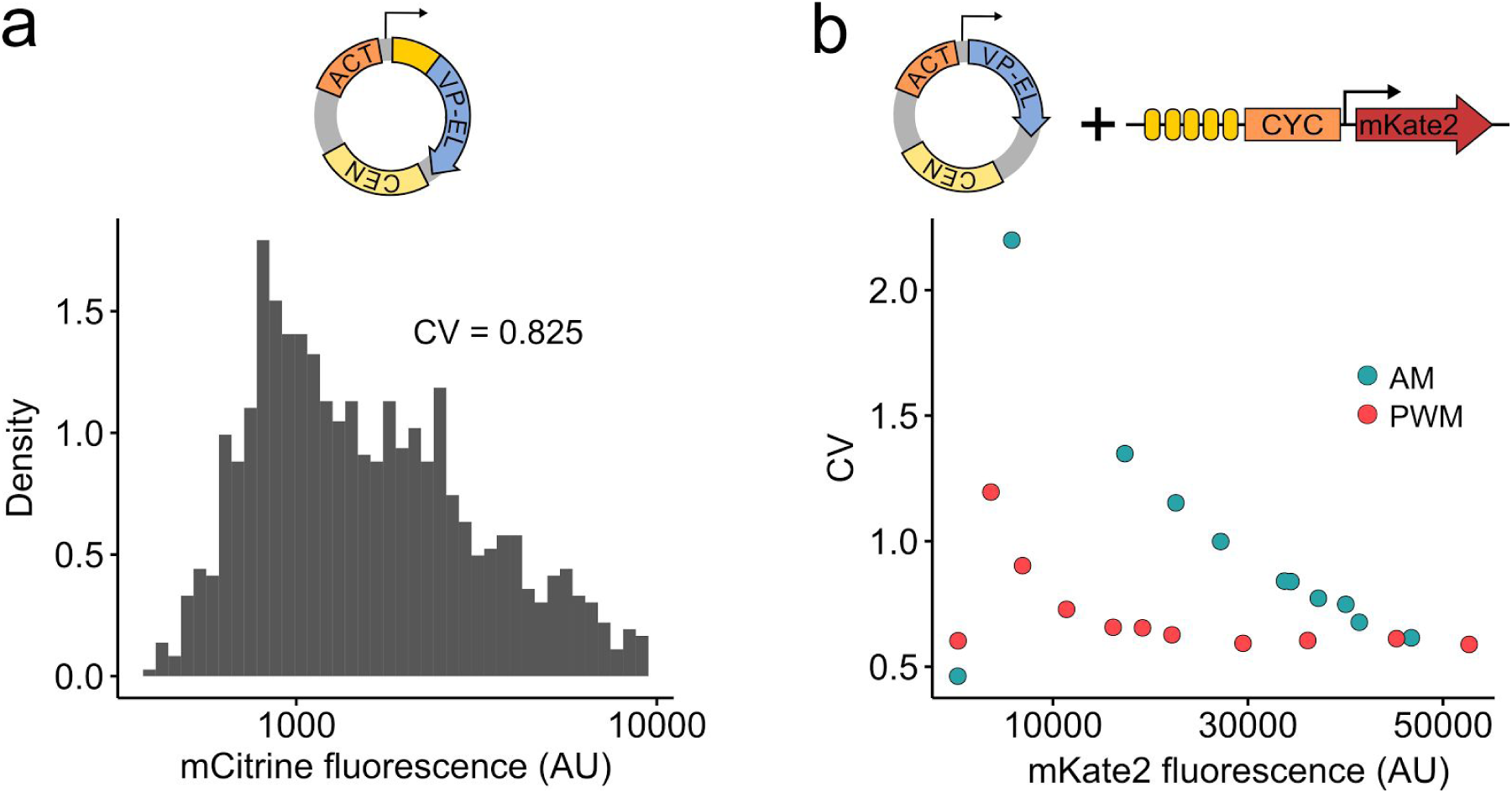
Effects of VP-EL222 expression from centromeric plasmid. (**a**) Distribution of mCitrine fluorescence as measured by flow cytometry. Cells express mCitrine-VP-EL222 under control of the ACT1 promoter from a centromeric plasmid (DBY128). mCitrine fluorescence was not normalized due to a low correlation of fluorescence with SSC. The CV of the fluorescence distribution is shown on the graph. (**b**) The CV is plotted as a function of mean expression from 5xBS-CYC180pr in response to AM (blue) and PWM (red) in a strain expressing VP-EL222 from a centromeric plasmid (DBY112). Cells were induced for 6 h before analysis. The following induction conditions were used: AM: 0, 35, 70, 105, 140, 175, 210, 245, 280, 350, 420 µW/cm^2^ light intensity. PWM: 420 µW/cm^2^ light intensity; 45 min period; 0, 3.3, 6.7, 13.3, 20, 26.7, 33.3, 46.7, 60, 90, 100 % duty cycle.

**Supplementary Fig. 12.**
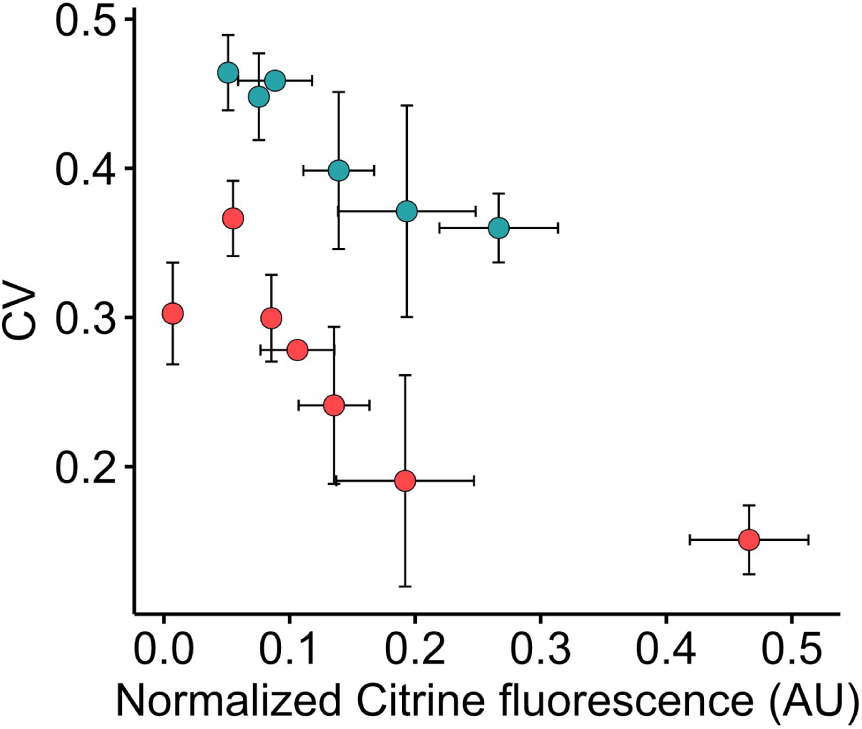
CV as a function of mean mCitrine expression regulated by AM (blue) and PWM (red) in a diploid strain expressing both Citrine and URA3 from 5xBS-CYC180pr (DBY125). Data is derived from the experiment shown in Fig 3g. Data represents the mean and s.d. of two independent experiments.

### Supplementary Note 1: Mathematical Modeling of the VP-EL222 mediated expression

In the following section, the specifics about the ordinary differential equation (ODE) model, parameter inference and model analysis are explained.

#### Suppl. Note 1.1 A simple ODE model describing VP-EL222 mediated gene expression

In order to obtain a quantitative understanding of the VP-EL222 based gene expression system in *S. cerevisiae*, we constructed a mathematical model of this process. The main purpose of the model is to describe the dynamics of the system in order to understand how dynamic inputs can be used to shape the gene expression output. We thus decided to employ a simplistic gene expression model consisting of three ODEs. These ODEs describe:

1. Light dependent activation and of the TF VP-EL222.
2. TF-dependent transcription and mRNA degradation.
3. Protein translation and degradation.

For simplicity, we assume that TF multimerization and promoter binding occur on fast timescales compared to the transcription process and we thus use Hill-type kinetics to model the effect of activated VP-EL222 (TF_on_) on the transcription rate.

The model is described by the following ODEs (I denotes the blue light input and TF_tot_ denotes the total amount of cellular VP-EL222):

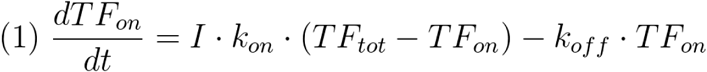

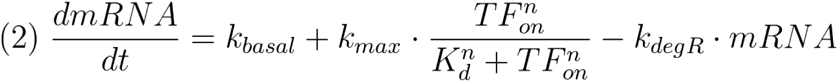

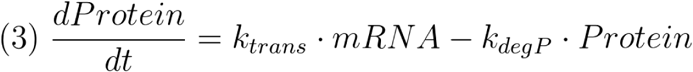

#### Suppl. Note 1.2 Fitting model parameters to experimental data

The model possesses 10 parameters (9 rate parameters and the total concentration of the TF / VP-EL222 (TF_tot_)). The value TF_tot_ acts as a scaling factor that can be by completely compensated for by changes in other parameters (k_on_ and K_d_) and does not affect the dynamics of the system. We thus fixed this value to 2000 molecules/cell. All characterization experiments were performed using the VP-EL222 mediated expression of the stable fluorescent protein (FP) mKate2. We thus equate the protein degradation rate (k_degP_) to the cellular growth rate of 0.007 min^-1^ (results of growth rate measurements are shown in **Supplementary Fig. 1a**). Thus, we end up with 8 free parameters that need to be estimated.

For this purpose, we performed three characterization experiments using the strain DBY43, expressing mKate2 from the 5xBS-CYC180 promoter.

1. We performed time-course measurements of mKate2 expression under constant illumination conditions. This experiment was performed to elucidate the kinetics of VP-EL222 activation and (to a larger extent) that of mRNA accumulation / degradation. The kinetics of protein accumulation are given by having fixed k_degP_.
2. We analyzed the dependence of mKate2 expression on light intensity, i.e. AM. This experiment gives us information about the mapping of light intensity to active transcription factor and finally transcription / protein expression.
3. We analyzed the dependence of mKate2 expression on the duty cycle in a PWM experiment with a short, 7.5 min period. The rationale behind this experiment is that it provides us with information about the kinetics of VP-EL222 activation and deactivation.

The results of these experiments are shown in **Supplementary Fig. 2**. We note again, that we are mainly interested in the dynamics of the gene expression system and not the absolute values of cellular mRNA or protein contents. For the model fitting, we thus assume a direct relation between fluorescence measured by flow cytometry and protein expression in the model.

Parameters were estimated by fitting the model to the mean of three independent experiments of each class of characterization experiments. To do so, we used a simplex-based search (Nelder-Mead algorithm, “fminsearch” function in Matlab) to minimize the sum of squared residuals (SSR) between the model and the data. This procedure was performed for different initial parameter values. The parameters resulting in the minimal SSR between all runs were used in this study and are reported in **Supplementary Table 1**. The model fits are shown in **Supplementary Fig. 2**.

#### Suppl. Note 1.3 Using the model to analyze functional regimes for PWM

The goal of PWM in this study is to regulate TF activity in a pulsatile fashion, while leading to close to constant protein levels over time at steady state. We used the mathematical model to analyze how these properties are affected by different parameters, mainly k_off_, k_degP_, and the PWM period. In order to ensure that we are not analyzing transient model behavior, all metrics described below are calculated after running the model for a simulated time of 720 min.

Effects of pulsatile TF regulation via PWM can be expected to be most pronounced when the concentration of active TF directly follows the light input, meaning that cellular TF activity itself shows either the maximal desired value or its basal level at any given time. However, in every realistic scenario, the temporal TF activity will deviate from this behavior to an extent that depends on the kinetics of TF activation / deactivation as well as the PWM period. We hence analyzed how the PWM period and the TF deactivation rate (k_off_) affect TF pulsing. In order to quantify this behavior, we use a tracking score defined by the ratio between the integrated TF activity during the light pulse and the whole period (**Fig. 1f**). This metric is 1 if the TF activity perfectly tracks the input and equals the duty-cycle if TF activity does not change over the PWM period. We calculated values of this metric for a duty cycle of 50% (**Fig. 1f**) As expected, the model shows that longer PWM periods are required with decreasing k_off_ to achieve a similar tracking score. The model further shows that the inferred rate of k_off_ for VP-EL222 (0.34 min^-1^, equivalent to a on-state half-life of about 2 min) is sufficiently large for performing PWM with reasonable periods. For a period of 30 min cellular TF activity is predicted to be at its maximal level during much of the light pulse and to return to basal levels in the dark before the next pulse (**Fig. 1f**). In contrast, when the period is reduced to 7.5 min, TF activity is predicted to reach its maximal activity before the end of the light pulse and to not return to the basal level in the dark (**Fig. 1f**). This leads effectively to gene regulation via mixed contributions of constant TF activity and weak pulsing. We found experimentally that this difference has strong functional consequences for the ability to use PWM for gene co-regulation (**Fig. 2d, Supplementary Fig. 7**) and noise reduction (**Fig. 3a**,**e**).

The model shows that pulsatile TF regulation can be more easily achieved with long PWM periods. However, using long PWM periods to regulate gene expression can potentially result in significant temporal fluctuation on the protein level, which is often not desirable. We thus sought to quantify the temporal response of protein expression to PWM. We use a score defined by the ratio of the maximal expression difference during the period divided by the mean expression level (**Fig. 1g**). Using this score, we found that even for a 30 min period, temporal changes in protein expression at steady state are expected to be minor for a wide range of protein half-lives (**Fig 1g**). We confirmed experimentally that there is no measurable input tracking for a stable fluorescent protein (**Fig 1g**). For the median protein half-life of ≈40 min in *S. cerevisiae* ^1^, PWM is predicted to lead to a maximal temporal fluctuation of about 6% for a 10% duty cycle. We note that this value is also affected by the mRNA degradation rate. Parameter estimation resulted in a value of 16.5 min for the mRNA half-life, which is close to the median half-life in S. cerevisiae (10 - 20 min) ^2,3^. Thus, modeling suggests that VP-EL222 should enable PWM-based regulation of a large percentage of yeast proteins. For short-lived proteins, the system would need to be optimized by introducing mutations that increase the dark-reversion rate. Such mutations were identified previously ^4,5^. We further note that VP-EL222 was previously employed in higher eukaryotes ^6^, where both mRNA and protein degradation rates were measured to be significantly lower than in *S. cerevisiae* ^7,8^. It is thus likely that PWM can be very successfully applied in these organisms and that PWM should also be possible with other tools for gene expression regulation that work on longer time-scales.

#### Suppl. Note 1.4 Estimating nascent RNA accumulation

Single-molecule FISH (smFISH) allows for the quantification of nascent transcripts, which is a fast readout of VP-EL222’s transcriptional activity. We performed an smFISH experiment in which we measure the transcriptional response of 5xBS-CYC180pr to a 20 min light pulse (**Supplementary Fig. 3**). To evaluate whether the identified model parameters describing

VP-EL222 activity and the transcription process are consistent with this data, we introduce an ODE describing nascent RNA accumulation:

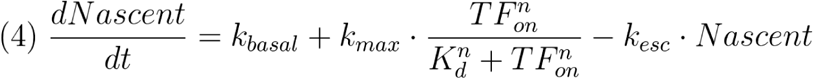

Here, TF-dependent nascent RNA production is modeled using Hill-type kinetics with the same parametrization as for mRNA production (**Equation 2, Supplementary Table 4**). The rate at which a nascent RNA escapes from the transcription site (k_esc_) is given by the RNA dwell-time which includes elongation and termination, leading to k_esc_ = (elongation time)^-1^ + (termination time)^-1^. The termination time was set to the literature value of 70 seconds ^9^ and the elongation time was set to 100 seconds based on the transcript length of 2000 bases and an average elongation rate of 20 bases per second ^9^. We found that the predicted dynamics of nascent RNA accumulation closely resemble the experimental data (**Supplementary Fig. 3c**). We note that this model is very simplistic - it does for example assume that nascent RNAs are observable (via smFISH) directly after transcription initiation.

#### Suppl. Note 1.5 Refitting of promoter-specific model parameters

In order to describe protein expression from the 2xBS-CYC180 promoter with the mathematical model identified above, we need to re-fit the promoter-specific parameters, k_basal_, k_max_, K_d_, and n. To do so, we performed two characterization experiment for this promoter, namely we measured the expression response to AM and PWM with a 7.5 min period. Model fitting was performed as described in **Suppl. Note 1.2**. Experimental results and fits are shown in **Supplementary Fig. 5**.

#### Suppl. Note 1.6 Modeling the effects of VP-EL222 variability on gene expression

Previous studies suggest that two major sources of extrinsic gene expression variability in *S. cerevisiae* are heterogeneity in TF expression and the cell cycle ^10,11^. Due to the fact that we directly affect TF dynamics, we thought to introduce cell-to-cell variability in TF concentration to our model and analyze the resulting CV-mean relationship for AM and PWM. As performed in other studies ^12^, we modeled protein / TF variability by running 10,000 ODE simulations differing only in the value for TF_tot_ for each input condition (consequences of this modeling choice are described below). Here, each simulation represents a single cell.

To do so, we first measured the fluorescence distribution of mCitrine tagged VP-EL222 to estimate heterogeneity of TF expression. We find that this distribution can be well described by a log-normal distribution with a CV of roughly 0.2 (**Supplementary Fig. 10a**). We then drew values for TF_tot_ from a log-normal distribution with a CV of 0.2 and a mean value of 2000 (the TF_tot_ value used for parameter estimation) and ran ODE simulations for a simulated time of 360 min. We ran simulations for different types of inputs (AM, and PWM with a 7.5, 15, and 30 min period) and different promoters (5xBS-CYC180pr and 2xBS-CYC180pr). Results of these simulations are shown in **Fig. 3e** and **Supplementary Fig. 10b.**

We found that the model can qualitatively recapitulate the CV-mean relationship for gene expression regulated by both AM and PWM. It also recapitulates the tunability of gene expression variability by changes in PWM period. In addition, the model predicts a reduced noise attenuation by PWM for the 2xBS-CYC180pr compared to the 5xBS-CYC180pr, which is verified by our experimental results (**Supplementary Fig. 10b**,**c**). This result shows the importance of working with “promoter-saturating-inputs” for maximal noise reduction by PWM (see **Fig. 3d** for an illustration and **Fig. 2b** for input-output functions for both promoters).

However, quantitatively, the model overestimated cell-to-cell variability in gene expression. This is likely a consequence of the model assumption that the TF concentration is fixed in each cell over the 6 h experiment. Indeed, when we compare the simulation results to measurements taken after 2 h of induction (**Supplementary Fig. 10d**), we find a better quantitative agreement. Furthermore, the model does not take into account intrinsic variability which is non-negligible at lower induction levels (**Fig. 3b**,**c**). We thus expect quantitative and qualitative differences between the model and the data for low expression levels, which can be seen in **Supplementary Fig. 10b**,**c**.

### Supplementary Note 2: DNA sequences

Details about the DNA constructs used in this study are described below. The color coding of the sequences corresponds to colors used in the preceding text.

### VP-EL222 constructs

#### pDB58; pDB116 / ACT1pr - VP-EL222 - CYC1term

pDB58 was used to construct yeast strains expressing VP-EL222 from the LEU2 locus. It consists of the **ACT1 promoter**, the coding sequence for **NLS-VP16-EL222** (derived from pVP-EL222 ^6^) and the **CYC1 terminator** integrated into an integrative vector based on the pRS vector series. The same construct was integrated into the centromeric plasmid pRG215 ^13^ to generate pDB116.

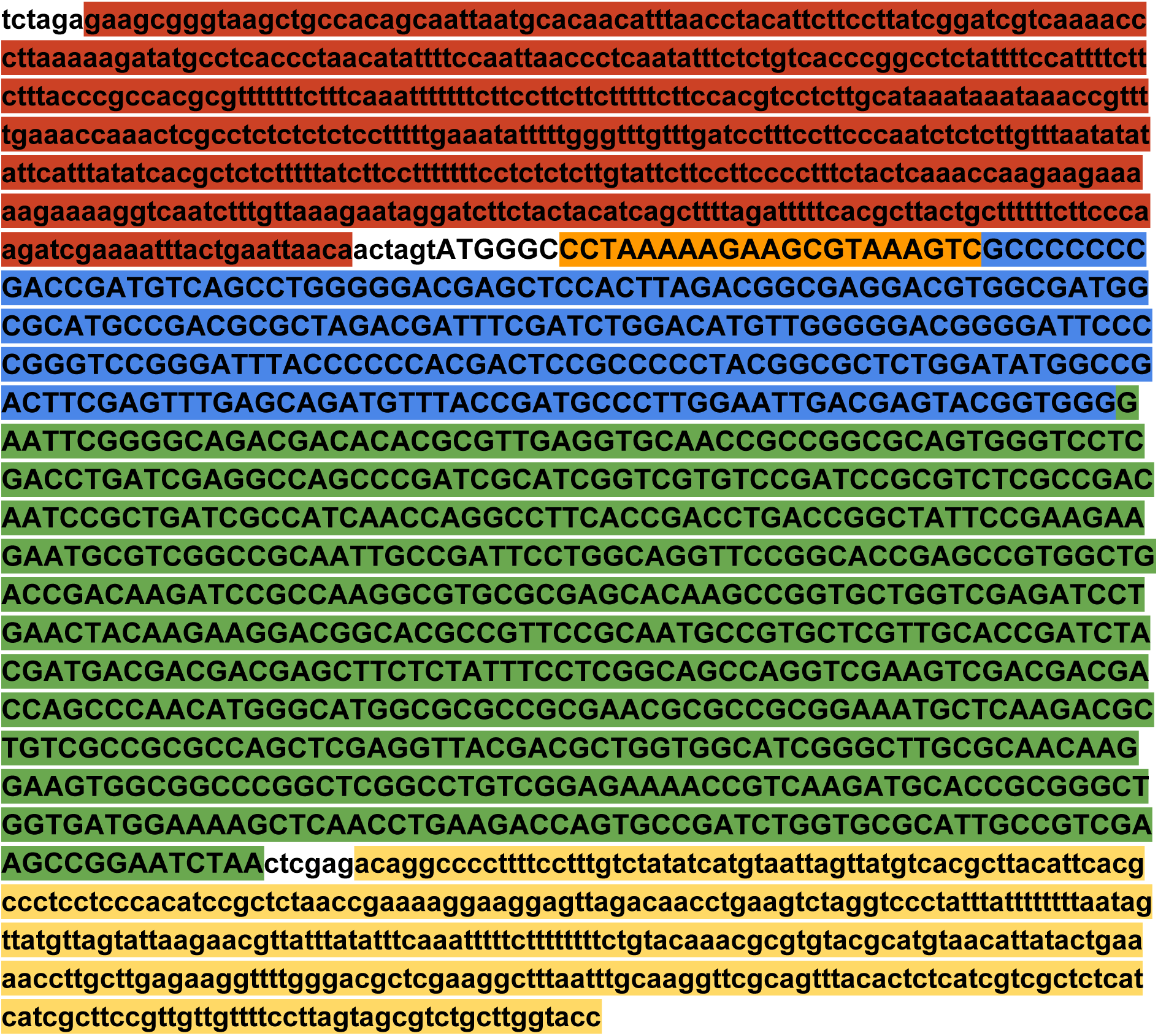

#### pDB113; pDB131 / mCitrine-VP-EL222

In order to quantify VP-EL222 expression in single cells, a coding sequence for **mCitrine** ^14^ was inserted upstream of VP-EL222. All other aspects of the sequence are as described above for pDB58 / pDB116

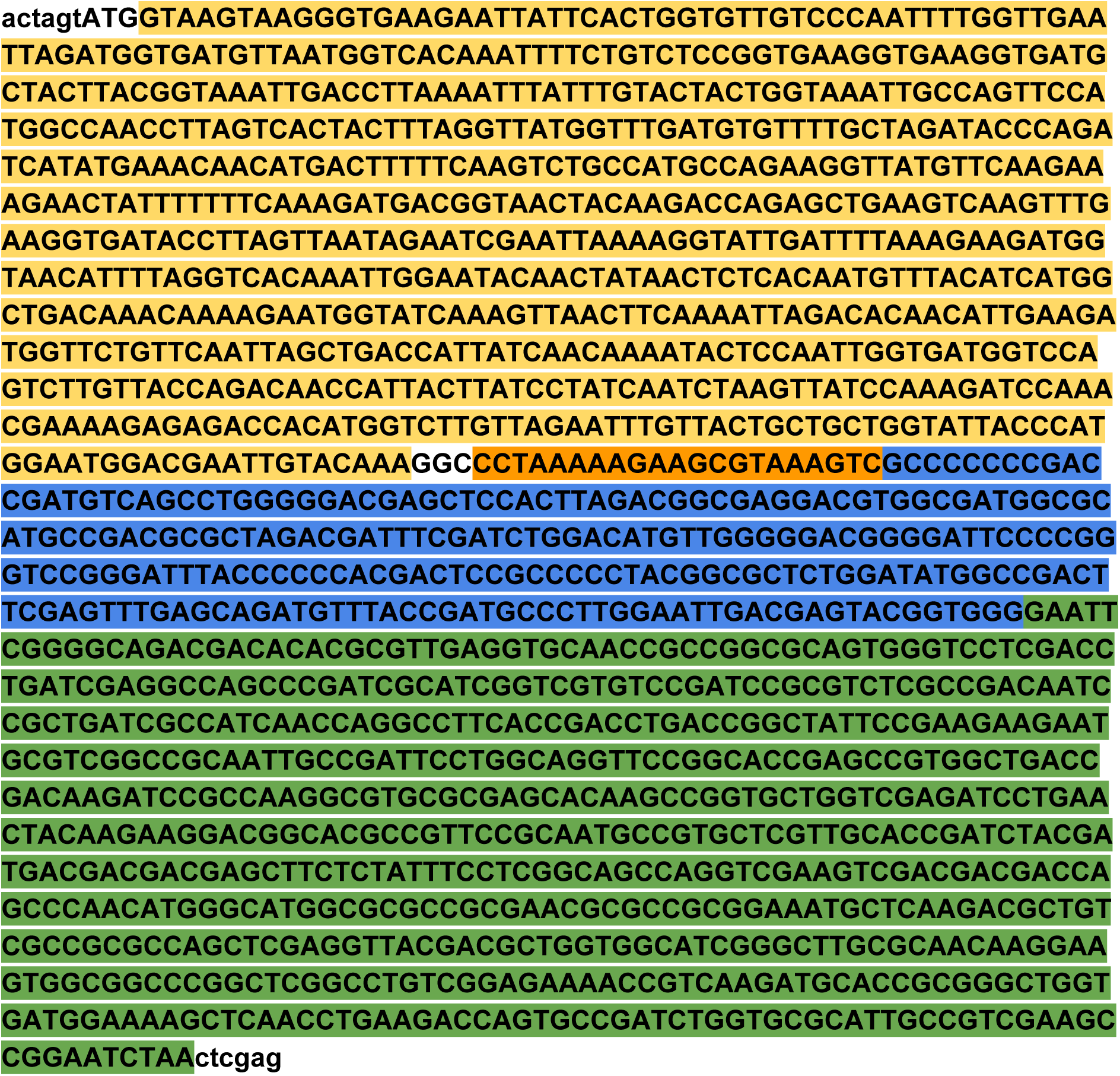

### VP-EL222 - dependent promoter sequences / reporter constructs

For all following promoter sequences, EL222 binding sites (BS; called C120 in the original publication) are underlined and promoter backbones are green. A sequence containing 5 binding sites for EL222 was amplified from pcDNA-C120-mCherry ^6^. All other binding site combinations were constructed by oligonucleotide annealing to obtain a single plasmid containing a single EL222 binding site, followed by duplications of this sequence using restriction enzyme cloning.

#### pDB60 / 5xBS-CYC180pr-Kozak-mKate2-ADH1t

pDB60 is used to express **mKate2** ^15^ under control of the 5xBS-CYC180 promoter. The promoter consists of a sequence containing 5 EL222 binding sites as well as a 180 bp sequence derived from the CYC1 promoter (**CYC180**). CYC180 was amplified from BY4741 genomic DNA ^16^. A consensus **Kozak** sequence was inserted upstream of the start codon to enhance translation. The mKate2 reporter gene is inserted into pFA6a-His3MX6 ^17^ using PacI and AscI sites (upstream of the **ADH1 terminator**). 5xBS-CYC180pr is inserted using HindIII and PacI.

All other VP-EL222 dependent promoters (see below) were characterized using the same plasmid backbone and were integrated into HindIII/PacI digested plasmid.

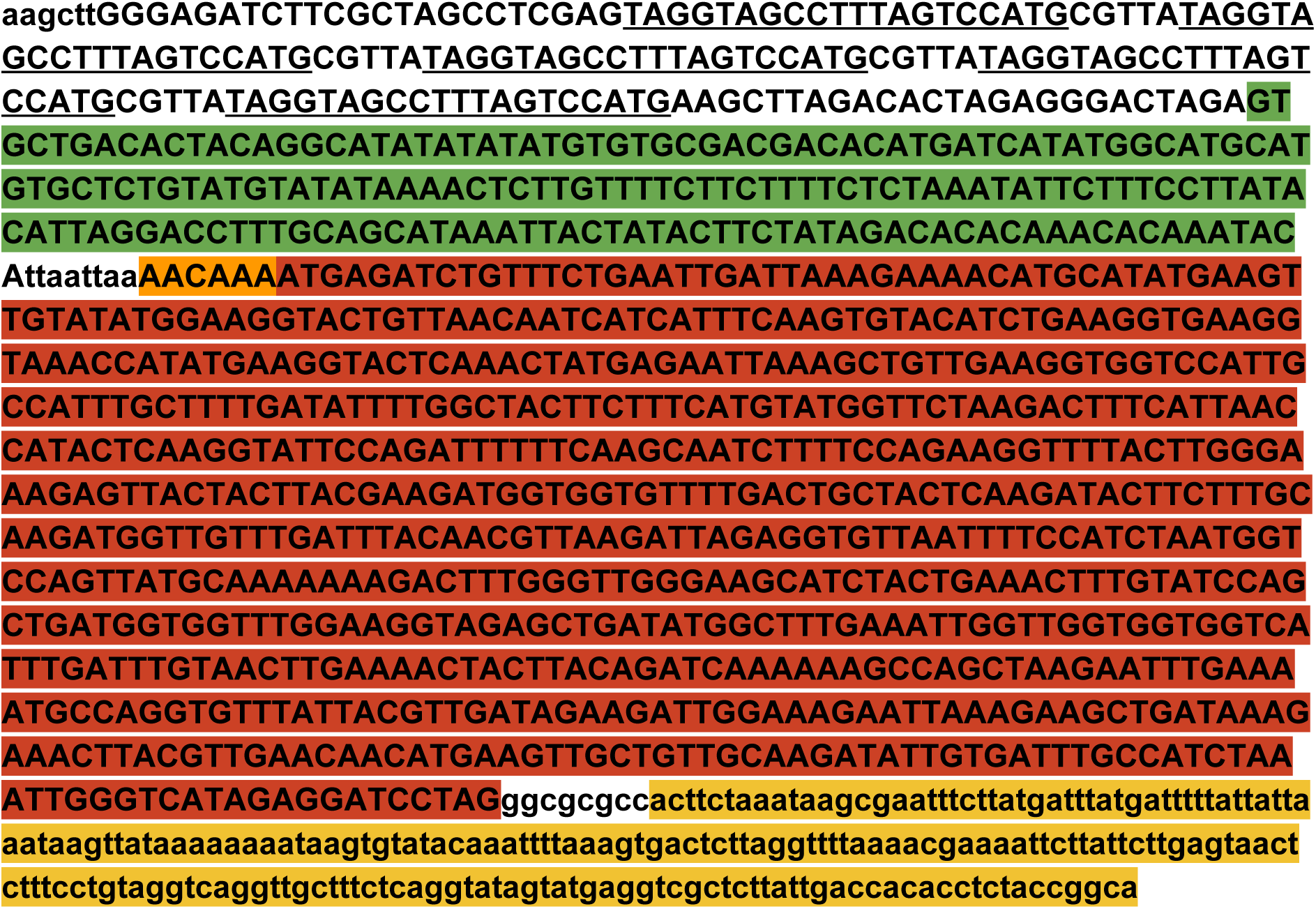

#### pDB72 / 5xBS-GAL1pr

The design of the VP-EL222 dependent GAL1-based promoter is adapted from another synthetic gene expression system presented in Ref. ^18^. The promoter was constructed by exchanging the UAS-GAL region (containing Gal4p binding sites) of the GAL1 promoter with the 5xBS sequence from pcDNA-C120-mCherry ^6^.

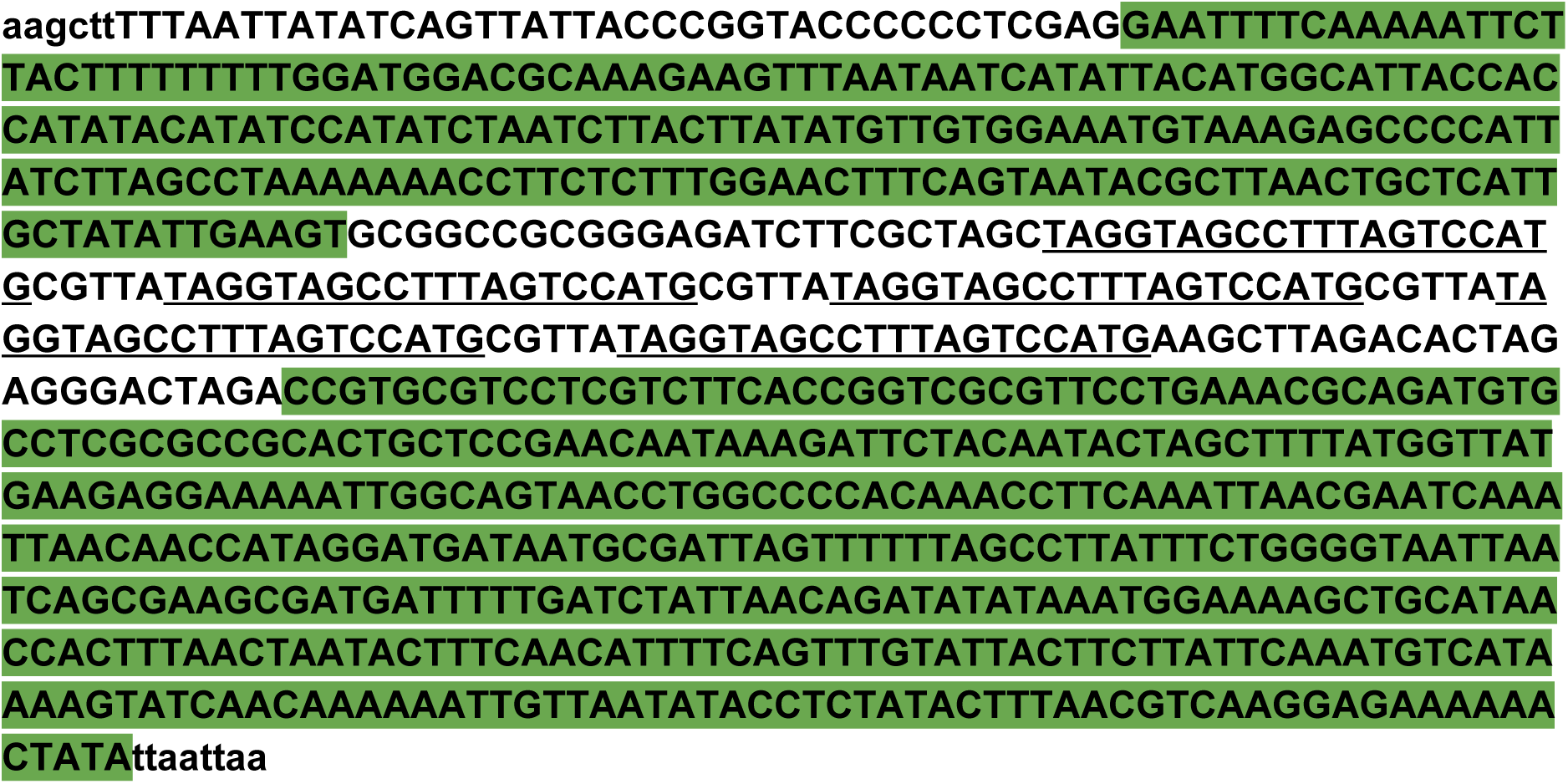

#### pDB107 / 5xBS-SPO13pr

In order to achieve light-dependant gene expression with very low basal expression, we inserted EL222 binding sites upstream of the basal SPO13 promoter ^19^. The SPO13 promoter sequence was amplified from BY4741 genomic DNA.

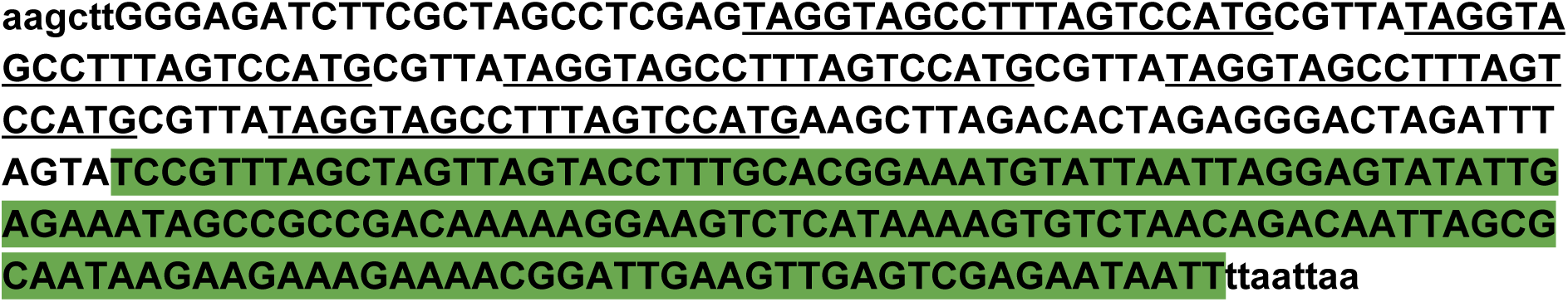

#### pDB99 / 2xBS-CYC180pr

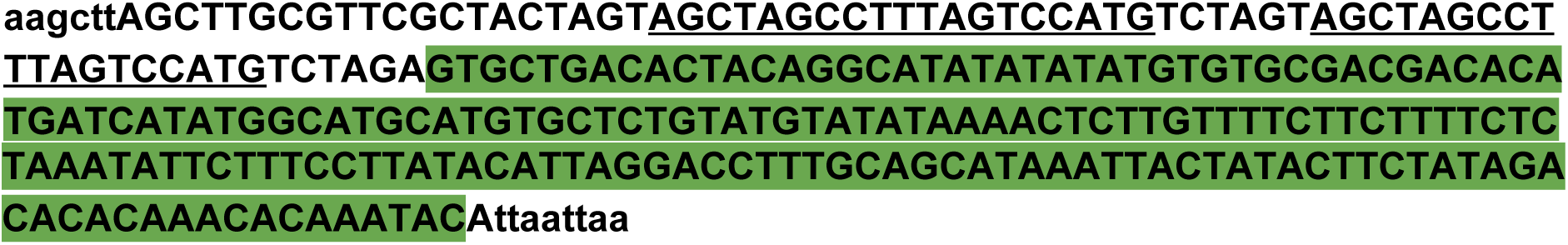

#### pDB100 / 3xBS-CYC180pr

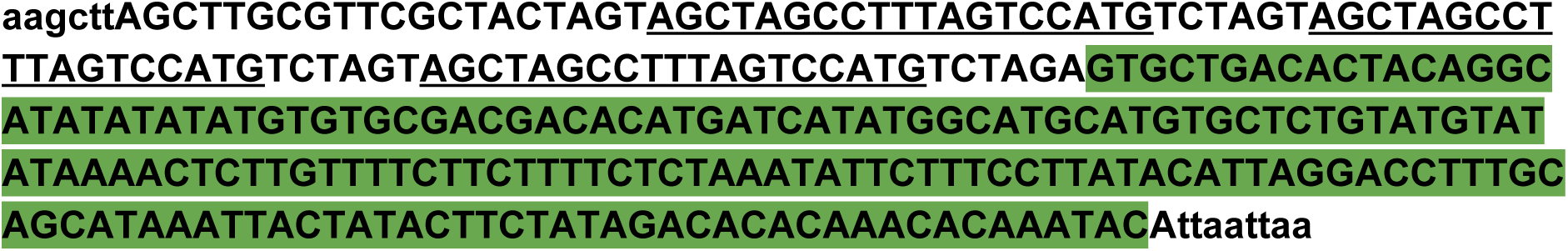

#### pDB101 / 4xBS-CYC180pr

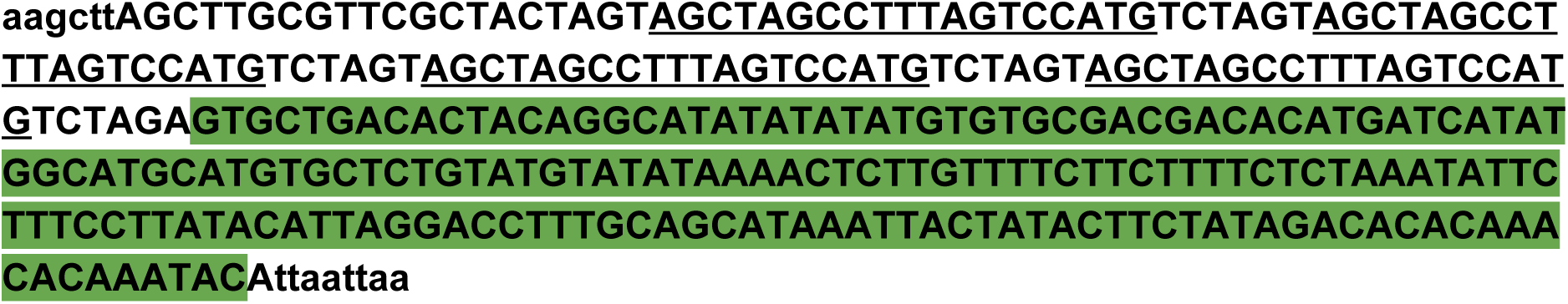

#### pDB102 / 6xBS-CYC180pr

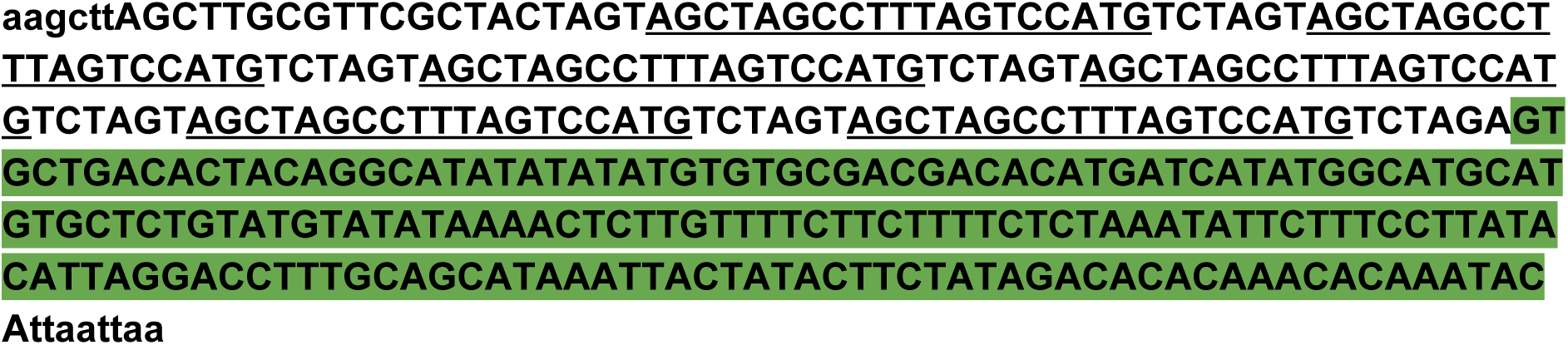

#### pDB103 / TDH3pr

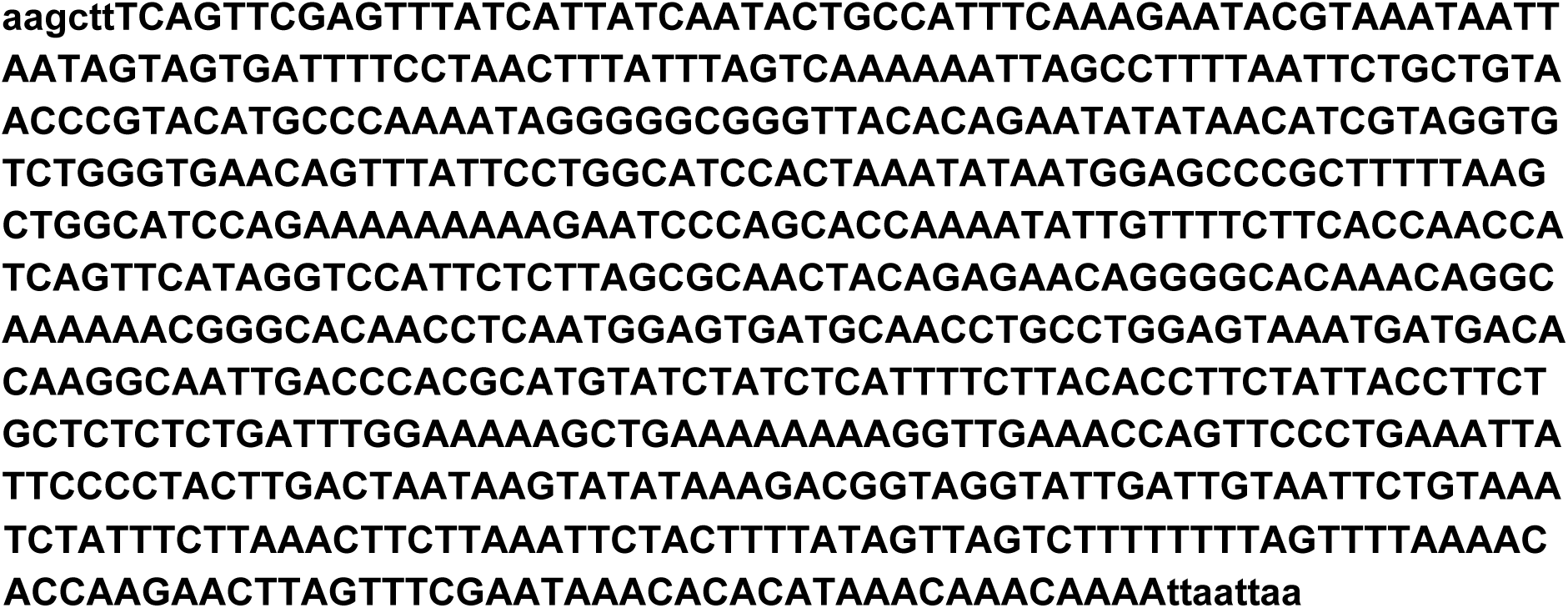

### Further sequences expressed from VP-EL222 dependent promoters

As shown above for mKate2 in pDB60, all sequences were integrated into PacI, AscI digested pFA6a-His3MX6-derived plasmids. Furthermore, all sequences possess a Kozak consensus sequence directly upstream of the start codon.

#### pDB78 / mKate2 - 24xPP7SL

A sequence containing 24 tandem repeats of the **PP7 stem loop** was amplified from pDZ416 ^20^ and was inserted after the mKate2 stop codon.

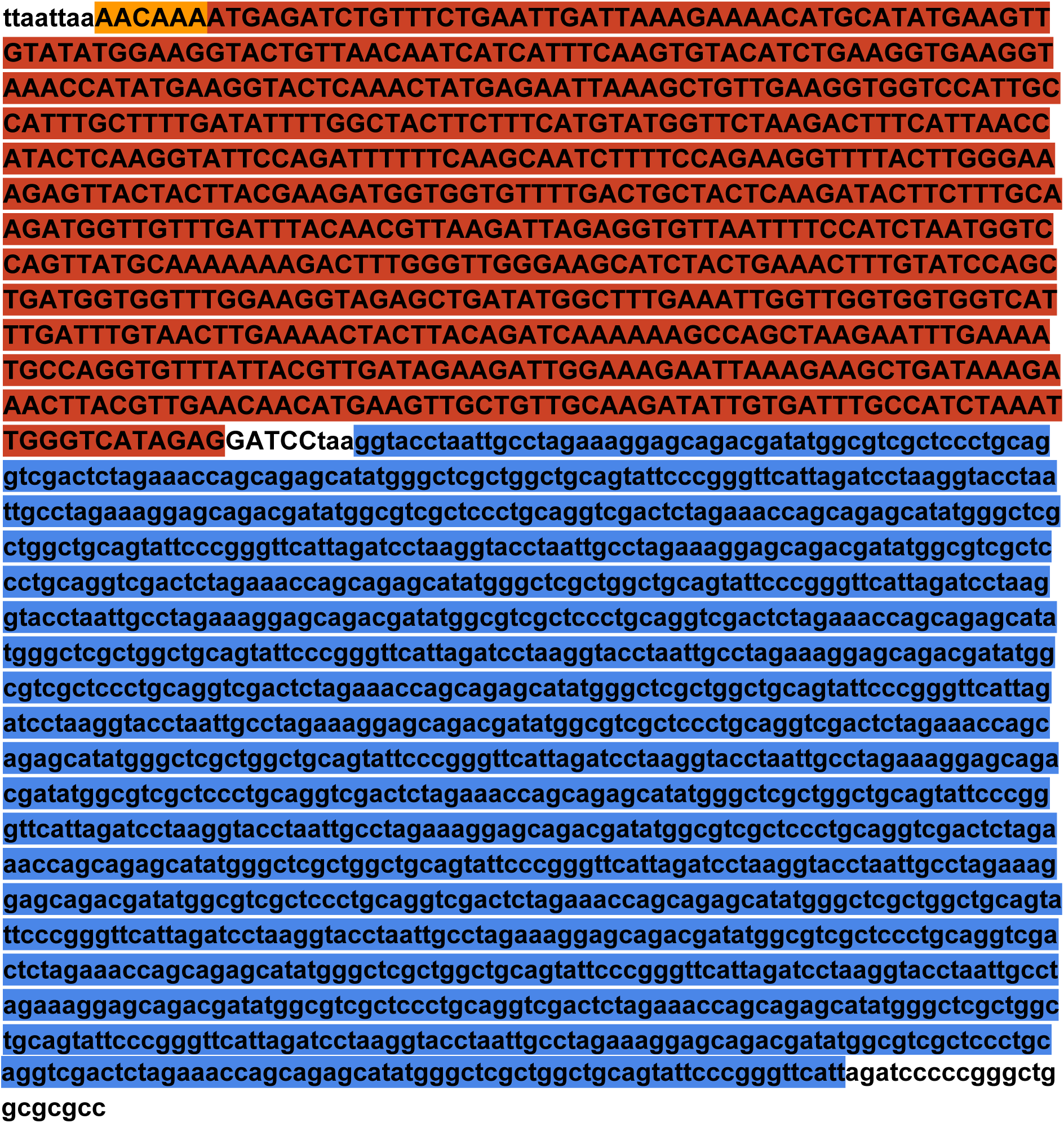

#### pDB110 / mCitrine

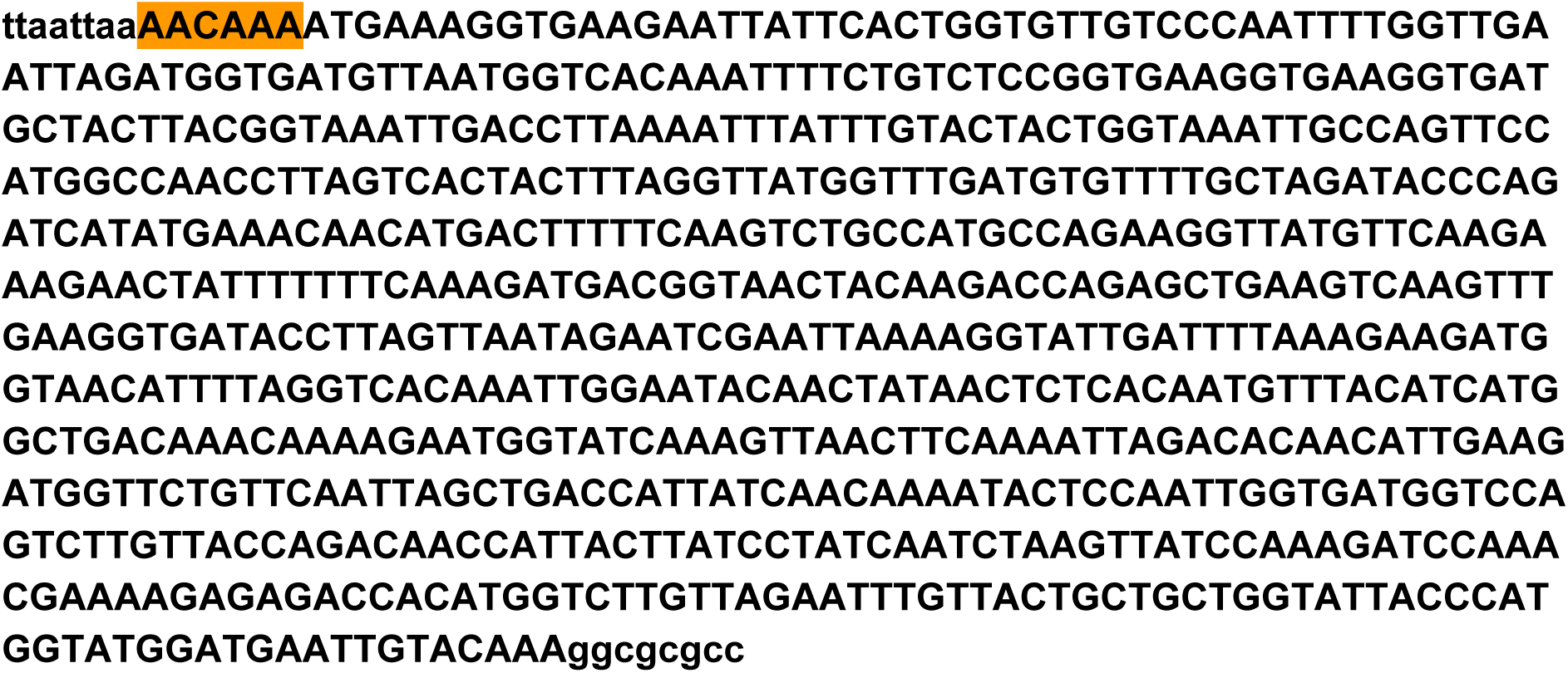

#### pDB111 / URA3

The URA3 coding sequence was amplified from BY4741 genomic DNA.

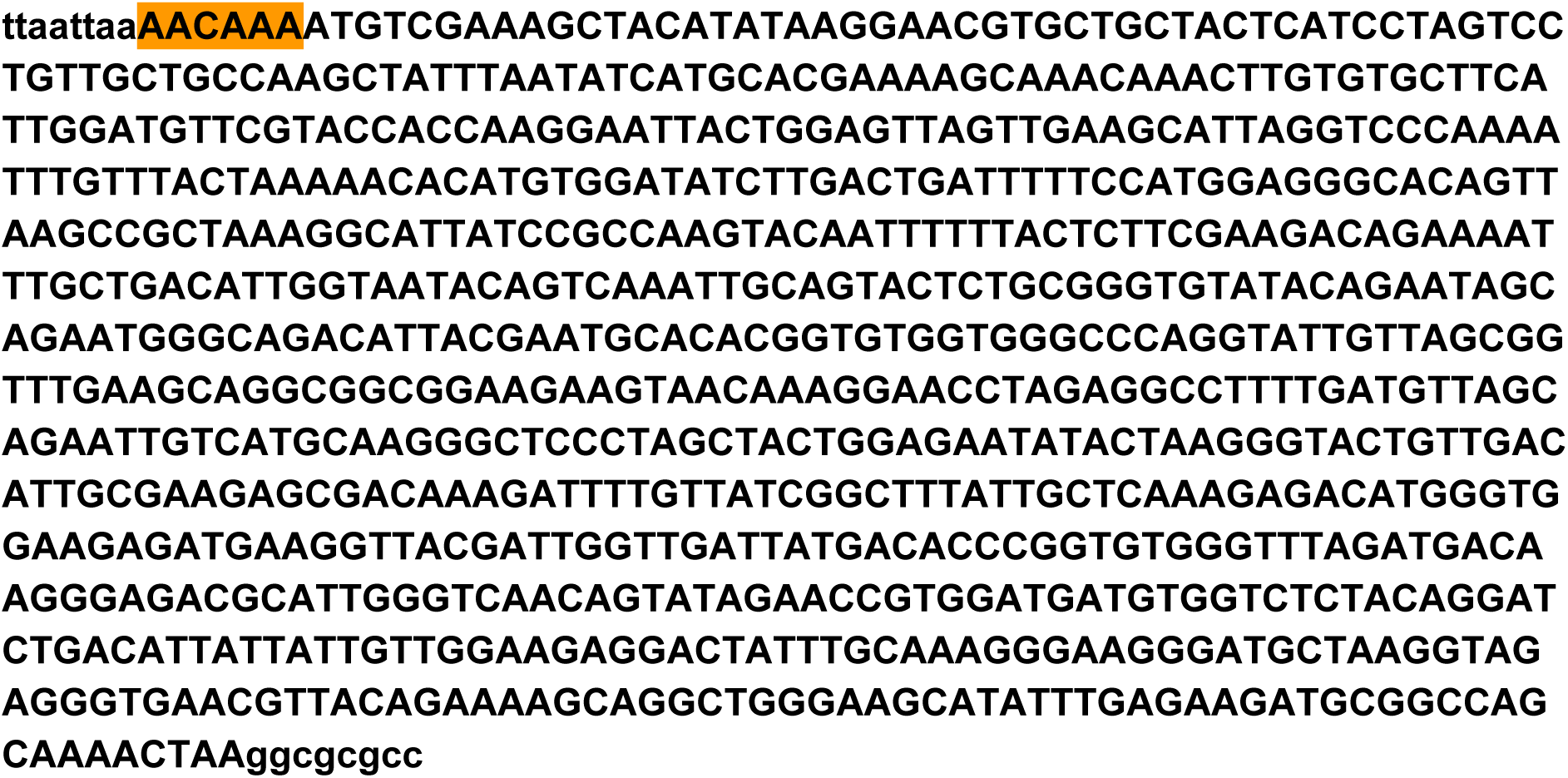

### Supplementary Tables

**Supplementary Table 1.** Plasmids used for strain construction. Promoters are represented by “pr”, terminators are represented by “t”.

| Plasmid | Backbone | Insert | Source |
| --- | --- | --- | --- |
| pDB58 | pKERG105 | ACT1pr-VPEL222-CYC1t | this study |
| pDB60 | pFA6-his3MX6 | 5xBS-CYC180pr-Kozak-mKate2-ADH1t | this study |
| pDB72 | pFA6-his3MX6 | 5xBS-GAL1pr-Kozak-mKate2-ADH1t | this study |
| pDB99 | pFA6-his3MX6 | 2xBS-CYC180pr-Kozak-mKate2-ADH1t | this study |
| pDB100 | pFA6-his3MX6 | 3xBS-CYC180pr-Kozak-mKate2-ADH1t | this study |
| pDB101 | pFA6-his3MX6 | 4xBS-CYC180pr-Kozak-mKate2-ADH1t | this study |
| pDB102 | pFA6-his3MX6 | 6xBS-CYC180pr-Kozak-mKate2-ADH1t | this study |
| pDB103 | pFA6-his3MX6 | TDH3pr-Kozak-mKate2-ADH1t | this study |
| pDB107 | pFA6-his3MX6 | 5xBS-SPO13pr-Kozak-mKate2-ADH1t | this study |
| pDB110 | pFA6-his3MX6 | 5xBS-CYC180pr-Kozak-Citrine-ADH1t | this study |
| pDB111 | pFA6-his3MX6 | 5xBS-CYC180pr-Kozak-KozURA3-ADH1t | this study |
| pDB113 | pKERG105 | ACT1pr-mCitrine-VPEL222-CYC1term | this study |
| pDB78 | pFA6-his3MX6 | 5xBS-CYC180pr-Kozak-mKate2-24xPP7SL-ADH1t | this study |
| pDB116 | pRG215 | ACT1pr-VPEL222-CYC1t | this study |
| pDB131 | pRG215 | ACT1pr-mCitrine-VPEL222-CYC1t | this study |

**Supplementary Table 2.** Strains used in this study. Promoters are represented by “pr”, terminators are represented by “t”.

| Name | Genotype | Source | Data shown in Figure |
| --- | --- | --- | --- |
| BY4741 | MATa his3 $\Delta$ 1 leu2 $\Delta$ 0 met15 $\Delta$ 0 ura3 $\Delta$ 0 | Euroscarf | 1d; S1a |
| BY4742 | MATalpha his3 $\Delta$ 1 leu2 $\Delta$ 0 lys2 $\Delta$ 0 ura3 $\Delta$ 0 | Euroscarf | - |
| DBY41 | BY4741, LEU2::ACT1pr-VPEL222-CYC1t(pDB57) | this work | 1d; S1a; S4 |
| DBY42 | BY4742, LEU2::ACT1pr-VPEL222-CYC1t(pDB57) | this work | - |
| DBY43 | DBY41, his3 $\Delta$ ::5xBS-CYC180pr-Kozak-mKate2-ADH1t-HIS3MX(pDB60) | this work | 1d,g; 2b-d, 3a, S1a, S4, S6, S8, S9 |
| DBY44 | DBY41, his3 $\Delta$ ::5xBS-GAL1pr-Kozak-mKate2-ADH1t-HIS3MX(pDB72) | this work | 2a; S4 |
| DBY69 | DBY41, his3 $\Delta$ ::2xBS-CYC180pr-Kozak-mKate2-ADH1t-HIS3MX(pDB99) | this work | 2a-d; S4; S5;S6 |
| DBY70 | DBY41, his3 $\Delta$ ::3xBS-CYC180pr-Kozak-mKate2-ADH1t-HIS3MX(pDB100) | this work | 2a; S4 |
| DBY71 | DBY41, his3 $\Delta$ ::4xBS-CYC180pr-Kozak-mKate2-ADH1t-HIS3MX(pDB101) | this work | 2a; S4 |
| DBY72 | DBY41, his3 $\Delta$ ::6xBS-CYC180pr-Kozak-mKate2-ADH1t-HIS3MX(pDB102) | this work | 2a; S4 |
| DBY122 | DBY41, his3 $\Delta$ ::5xBS-SPO13pr-Kozak-mKate2 -ADH1t-HIS3MX(pDB107) | this work | 2a; S4 |
| DBY88 | DBY41, his3 $\Delta$ ::5xBS-CYC180pr-Kozak-mKate2-24xPP7SL-ADH1t-HIS3MX(pDB78) | this work | S3 |
| DBY73 | BY4741, his3 $\Delta$ ::5xBS-CYC180pr-Kozak-mKate2-ADH1t-HIS3MX(pDB60) | this work | 1d |
| DBY100 | DBY41, his3 $\Delta$ ::TDH3pr-Kozak-mKate2-HIS3MX(pDB103) | this work | S1b; S8a,b |
| DBY104 | DBY42, his3 $\Delta$ ::5xBS-CYC180pr-Kozak-Citrine-ADH1t-HIS3MX(pDB110) | this work | - |
| DBY105 | DBY73, LEU2::pAct1-mCitrine-VPEL222-CYC1t(pDB113) | this work | 3f; S10 |
| DBY110 | MATa/MATalpha, DBY43/DBY104 | this work | 3b,c; S9 |
| DBY112 | DBY73, Act1pr-VPEL222-CYC1t (pDB116) | this work | 3g; S11 |
| DBY118 | MATa/MATalpha, DBY69/DBY104 | this work | S7 |
| DBY123 | DBY41, his3 $\Delta$ ::5xBS-CYC180pr-Kozak-KozURA3-ADH1t-HIS3MX(pDB111) | this work | - |
| DBY125 | MATa/MATalpha, DBY123/DBY104 | this work | 3h; S12 |
| DBY128 | DBY73, ACT1pr-mCitrine-VPEL222-CYC1t (pDB131) | this work | S11 |

**Supplementary Table 3.**
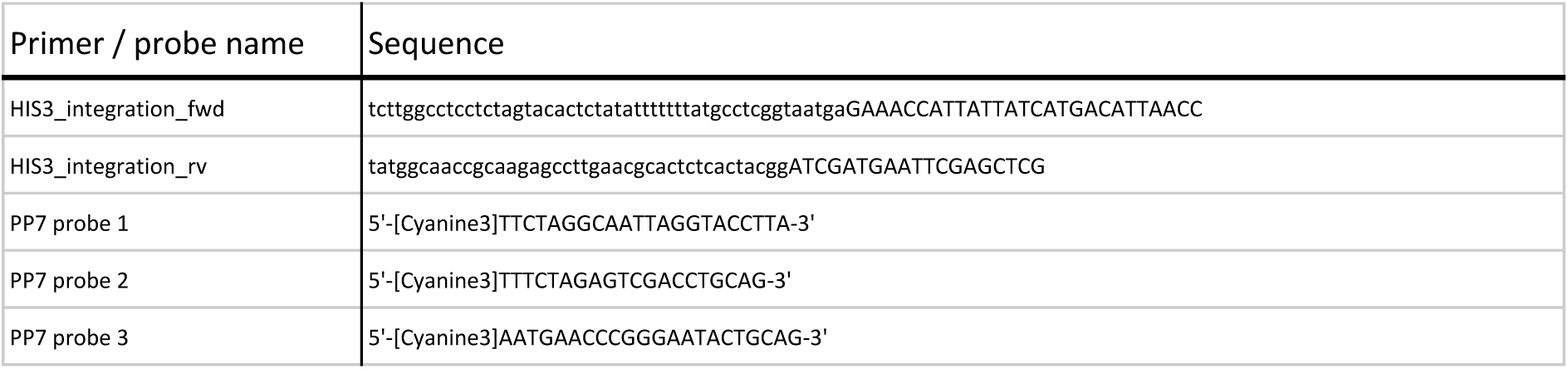
Primers and smFISH probes used in this study. For the HIS3-integration primers, uppercase bases are complementary to plasmid sequences and lowercase bases are complementary to the yeast genome. All smFISH probes are labeled with CY3 at the 5’ end. Probe sequences were obtained from Ref. ^4^.

**Supplementary Table 4.** Estimated parameters for the VP-EL222 model.

| Parameter | Description | value: 5xBS-C180pr | value: 2xBS-C180pr |
| --- | --- | --- | --- |
| $TF_{total}$ (molecules) | total cellular TF | 2000 | 2000 |
| $k_{on}$ ($\text{min}^{-1} * (\text{uW} / \text{cm}^2)^{-1}$) | light dependant VP-EL222 activation rate | 0.0016399 | 0.0016399 |
| $k_{off}$ ($\text{min}^{-1}$) | VP-EL222 dark-state reversion rate | 0.34393 | 0.34393 |
| $k_{basal}$ ($\text{mRNA} * \text{min}^{-1}$) | basal transcription rate | 0.02612 | 0.24358 |
| $k_{max}$ ($\text{mRNA} * \text{min}^{-1}$) | maximal induced transcription rate | 13.588 | 11.031 |
| $K_d$ (molecules) | $TF_{on}$ level required for achieving $k_{max} / 2$ | 956.75 | 1462.5 |
| $n$ (-) | hill coefficient | 4.203 | 4.6403 |
| $k_{degR}$ ($\text{min}^{-1}$) | mRNA degradation rate | 0.042116 | 0.042116 |
| $k_{trans}$ ($\text{proteins} * \text{min}^{-1} * \text{mRNA}^{-1}$) | translation rate | 1.4514 | 1.4514 |
| $k_{degP}$ ($\text{min}^{-1}$) | protein degradation rate | 0.007 | 0.007 |

**Supplementary Table 5.** Parameters for hill function fit describing the mapping of Ura3 expression to cell growth (shown in Fig. 3h). Equation: Growth-rate = k_b_ + k_m_ * Ura3^n^ / (Ura3^n^+ K_d_^n^).

| Parameter | AM | PWM |
| --- | --- | --- |
| $k_b$ ($\text{h}^{-1}$) | 0.0048 | 0.0062 |
| $k_m$ ($\text{h}^{-1}$) | 0.4034 | 0.4098 |
| $n$ | 2.7246 | 5.0183 |
| $K_d$ | 3292.8563 | 2936.0917 |

